# Long-range single-molecule mapping of chromatin modification in eukaryotes

**DOI:** 10.1101/2021.07.08.451578

**Authors:** Zhe Weng, Fengying Ruan, Weitian Chen, Zhe Xie, Yeming Xie, Chen Zhang, Zhichao Chen, Juan Wang, Yuxin Sun, Yitong Fang, Mei Guo, Yiqin Tong, Yaning Li, Chong Tang

## Abstract

The epigenetic modifications of histones are essential markers related to the development and pathogenesis of diseases, including human cancers. Mapping histone modification has emerged as the widely used tool for studying epigenetic regulation. However, existing approaches are limited by fragmentation and short-read sequencing represent the average chromatin status in samples and cannot provide information about the long-range chromatin states. We leveraged the advantage of long read sequencing to develop a method “BIND&MODIFY” for profiling the histone modification of individual DNA fibers. Our approach is based on the recombinant fused protein A-M.EcoGII, which tethers the methyltransferase M.EcoGII to the protein binding sites and locally labels the neighboring DNA regions through artificial methylations. We demonstrated that the aggregated BIND&MODIFY signal matches the bulk-level ChIP-seq and CUT&TAG, verify the single-molecule heterogenous histone modification status, and quantify the correlation between distal elements. This method could be an essential tool in future third-generation sequencing ages.

## Introduction

The genome in each cell of a multi-cellular organism is identical and stays reasonably static, whereas the epigenome is very dynamic ^1^. The epigenome varies in different cell types and plays significant roles in various biological processes, such as stem cell differentiation ^2^, the nervous system ^3^, cell aging and disease ^4^ . However, current methods developed to study the epigenome are vague and may be less comprehensive than those to study the genome, which can be directly sequenced. The solutions proposed have been based on extracting the epigenome signals by enzymatic or chemical means and isolating either the accessible or protected location. Therefore, much interest has been focused on collecting and comparing genome-wide chromatin accessibility and chromatin modification to locate the specific epigenetic changes that accompany cell differentiation, environmental signaling, and disease development ^5^. Next-generation sequencing using DNase I (DNase-seq) and MNase (MNase-seq) treatment has provided the first demonstration that active chromatin coincides with nuclease hypersensitivity ^6 7^. A comparable method is ATAC-seq which screens for the transposon hyperactivity on the accessible chromatin ^8, 9^. The most recent and advanced theory that proposed to offer a single molecular long-read approach for evaluating chromatin accessibility is a third-generation sequencing technique, referred to as nanopore sequencing, that includes SMACseq ^10^, NOME-seq ^11^, and nanoNOME ^12^. Chromatin accessibility assays may address certain epigenetic problems, but the more specific question “which protein regulates chromatin accessibility” cannot be answered. The dynamic chromatin structure is tightly regulated by post-translational histone modifications and binding transcription factors ^13^. To answer the question: “is a transcription factor bound to a piece of DNA or locate the histone modification” scientists use “ChIP-seq” to find out if a “protein of interest” is bound to a piece of DNA ^14^. Large-scale chromatin immunoprecipitation assay (ChIP-seq) utilize specific antibody to precipitate DNA fragments crosslinked with target proteins, followed by direct ultra-high-throughput DNA sequencing ^15^. ChIP-seq has facilitated the finding of protein-binding motifs and has allowed us to identify noncanonical protein-binding motifs further. In the literature, these have been extensively investigated to study the biological functions of the histone acetylation and methylations, transcription factors, etc. ^16–22^. Currently, the major challenge restricting the use of ChIP-seq is the need for high input DNA amounts. There have been certain methods established to conduct ChIP-seq with low input DNA, for example, ChIPmentation ^23^ CUT&RUN ^24^ and CUT&TAG ^25^. For the CUT&RUN protocol, MNase is fused to protein A (pA-MNase) to guide the chromatin cleavage to antibodies bound to the protein targets at their binding sites across the genome ^25^. Conceptually similar work has also been carried out by CUT&TAG, in which transposons are used instead of the MNase. Many of these techniques and their derivatives have been improved to achieve the single-cell DNA input ^25–28^. However, the complex relationship between DNA methylation, chromatin modification, and genome structure variation is often difficult to unravel with a single omics tool. Various methods developed to address this challenge involving applying the bisulfite treatment to the immunoprecipitated DNA fragments during ChIP-seq, which ensures that both DNA methylation information and histone information are procured from the process ^29–31^. The above literature review describes the previous and current attempts to address the status of the protein-bound on DNA.

Most ChIP-seq technologies employ the short-read sequencing by next-generation sequencing (NGS), and downstream analysis has been based on the peak calling algorithm with statistical analysis of populated fragments ^8, 9^. As a result, researchers are then faced with a question - whether the individual chromatin fibers retain the same long-range physical organization as they remove linkages between distal segments. Moreover, due to its use of short-read sequencing, the methylation and structure variation information is limited to the immunoprecipitated genome region. In addition, these short-read sequencing technologies do not utilize non-cleavage labeling methods and lack the capability of long-read sequencing to preserve all necessary epigenetic information in individual chromatin.

Our new approach BIND&MODIFY, by fusing methyltransferase to bind and modify the targeted sites within a long DNA molecule. Furthermore, we developed a single-molecule long-read bound protein mapping sequencing method. This single-molecule method directly examines the protein binding regions, DNA methylation, and complex structure within a single chromatin fiber at multikilobase scales. We used BIND&MODIFY to study histone methylation and co-methylated histone status in cancer cells. Moreover, we enumerate the detail regulation in the transposon region and bivalent regulatory states, while observing distance coordinated changes in cancer cells. BIND&MODIFY also allows for the footprinting of other specific transcription factors. We expect future applications of BIND&MODIFY to enable new insights into the status of the dynamic regulators of the genome in various experimental systems and sequencing platforms.

## Results

### The experimental overview of the BIND&MODIFY method

The rationale of the BIND&MODIFY method is to indirectly label DNA regions bound with protein of interest via an engineered recombinant fusion protein, protein A-M.EcoGII (pA-M.EcoGII), whose methyltransferase activity can be locally controlled. The recently characterized adenosine methyltransferase M.EcoGII was capable of methylating a broad range of genome DNA in a sequence non-specific manner^32^ compared with DamID^33^, which could only methylate the rare GATC motif in the genome (Supplemental Figure 1). Our BIND&MODIFY approach leverages the power of an engineered recombinant protein, pA-M.EcoGII. First, the recombinant methyltransferase was tethered close to the specific antibody-bound protein of interest, and the adenines in the DNA sequences adjacent to the protein of interest were methylated at m^6^A in a non-specific manner upon activation (Figure 1A). As m^6^A modifications are very limited in the native background of the mammalian chromosome ^34, 35^, the artificial m^6^A modification, indicating the protein binding motif, could be directly detected by the nanopore ^36, 37^.

**Figure 1.**
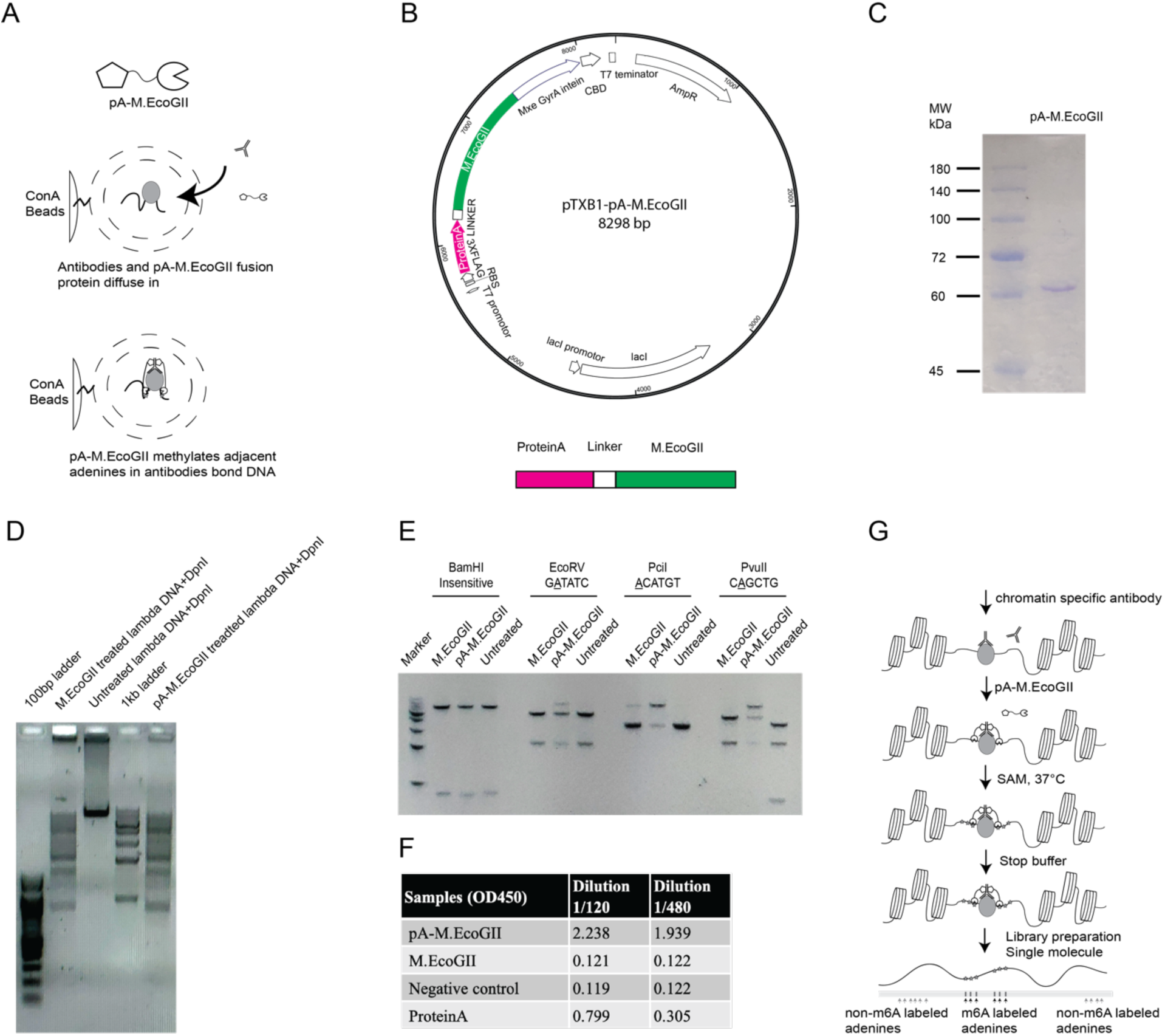
The experiment concept and validation of the recombinant protein pA-M.EcoGII in BIND&MODIFY. (A)The experiment concept of the BIND&MODIFY. The cells were lightly fixed and permeabilized for the antibody and enzyme passage. The recombinant protein pA-M.EcoGII was located to desired sites under antibody guidance. Then the M.EcoGII activity was locally activated to modify the nearby regions to label the genomic DNA with targeted binding proteins. (B) The upper panel showed the plasmid map of the pA-M.EcoGII. The fused pA-M.EcoGII was cloned into pTXB1 plasmid and purified with compatible IMPACT protein purification system. The lower panel showed the expressed fusion protein structure: Protein A-linker-M.EcoGII-intein-CBD. (C) The Coomassie blue gel stain showed the purity of the purified protein A-M.EcoGII. (D) Methylation of linear lambda DNA by pA-M.EcoGII activates m6A-site dependent DpnI restriction endonuclease digestion. The PCR amplified unmethylated lambda DNA was treated with commercial M.EcoGII, no enzyme, and pA-M.EcoGII. The G<u>A</u>TC m6A methylation dependent restriction endonuclease DpnI digestion showed the comparable methyltransferase activity of the commercial M.EcoGII and our recombinant proteins. DNA marker: 100bp ladder, 100-1510bp(left); 1kb ladder, 250-10,000bp(right). (E) Methylation of linear dsDNA by pA-M.EcoGII inhibits multiple site-specific methylation sensitive restriction endonucleases. The unmethylated DNA template was a 7kb linear dsDNA, which was PCR amplified from pTXB1 plasmid. The DNA template was treated with commercial M.EcoGII, pA-M.EcoGII and no enzyme. These treated DNA templates were each incubated with four restriction endonucleases (BamHI, EcoRV, PciI, PvuII). The BamHI is the m6A methylation insensitive enzyme, and the EcoRV, PciI, PvuII are the m6A methylation sensitive enzyme, with which the digestion could be blocked by corresponding m6A site. Our pA-M.EcoGII recombinant protein showed digestion inhibition on EcoRV, PciI, PvuII digested samples, better than commercial M.EcoGII, as compared to untreated DNA template. DNA marker, 1kb ladder, 250-10,000bp. (F) The antibody affinity assay showed the recombinant pA-M.EcoGII had the affinity to the secondary antibody in two different dilutions (1/120,1/480, 10mg/ml). (G) The experiment outlines of BIND&MODIFY. After light fixation and permeabilization, the cells were tethered to Concanavalin A magnetic beads for the purification in the next steps. Then the cells were incubated with antibody and pA-M.EcoGII with minimal washes. The addition of S-adenosylmethionine to initialize the methylation reaction. The DNA was extracted to prepare the library for ONT nanopore sequencing. After sequencing, the data was processed as genome alignment and m6A base calling.

Details of recombinant protein pA-M.EcoGII design, construction, and purification steps can be found in the methods section. Briefly, recombinant pA-M.EcoGII is designed by cloning two immunoglobin binding domains of staphylococcal protein A fused N-terminally with M.EcoGII (Figure 1B). The amino acid of the linker region between protein A and M.EcoGII is DDDKEF. The recombinant pA-M.EcoGII was expressed in the *E.coli* system and had a molecular weight of 61 kDa (Figure 1C). To access the function of purified recombinant pA-M.EcoGII, the m^6^A dependent restriction enzyme DpnI digestion was used to verify pA-M.EcoGII’s methylation activity on lambda DNAs. We found that pA-M.EcoGII processed similar m^6^A methylation specificity compared to commercially available M.EcoGII (NEB) (Figure 1D). To further evaluate the effectiveness of purified recombinant pA-M.EcoGII, m^6^A sensitive restriction enzymes (EcoRV, PciI, PvuII) were employed to cleave pTXB1 plasmid DNAs treated with pA-M.EcoGII and commercial M.EcoGII (NEB). Both pA-M.EcoGII and M.EcoGII successfully introduced the m^6^A methylations into the plasmid, which inhibited the digestion activity of m^6^A sensitive restriction enzymes (Figure 1E). We performed an ELISA assay to evaluate the potential of pA-M.EcoGII’s function of IgG domain binding and observed that it processed similar activity to commercially available protein A (Figure 1F). Taken together, our purified recombinant pA-M.EcoGII has comparable enzymatic activity to commercially available protein A and M.EcoGII.

Based on the recombinant pA-M.EcoGII protein, we developed the BIND&MODIFY protocol (Figure 1G). Briefly, the cells are 1) lightly fixed and tethered to Concanavalin A (ConA)-coated magnetic beads, 2) incubated with primary antibodies, 3) incubated with pA-M.EcoGII with minimal washes, 4) S-adenosylmethionine is added and incubated at 37°C for 30min to initialize the methylation reaction, and then quenched by 0.1% SDS, 5) extraction of DNA by phenol-chloroform and prepare the library for Oxford Nanopore Technology (ONT) sequencing. After sequencing, the data is processed through genome alignment and m^6^A signal detection. The m^6^A possibility cut-off was determined based on background noise (Supplemental Figure 2). The sensitivity and specificity of the m^6^A signal are around 0.92 and 0.99, respectively.

### The validation of BIND&MODIFY on assigned location *in vitro*

According to previous studies, the frequency of adenosine occurrence on various genomes is once in every 3bp ^10^, which was the theoretical resolution of M.EcoGII treated areas. The resolution of BIND&MODIFY depends not only on the adenosine frequency on the genome but also on the range of the tethered M.EcoGII could reach. Here we present the test that evaluates the signal resolution of BIND&MODIFY. The first step to developing the method was to synthesize the DNA with an assigned antibody binding site. Then, the PCR was used to introduce the m^5^C modification into the given location on DNAs (Figure 2A). In the following steps, the pA-M.EcoGII recombinant protein was tethered to the designed site with the help of the m^5^C antibody. Then the attached M.EcoGII methyltransferase transferred a methyl group to neighboring adenosine after activation. The N6-methyladenosine (m^6^A) modified DNAs were sequenced in a Nanopore device, and we performed in-house pipeline data analysis.

**Figure 2.**
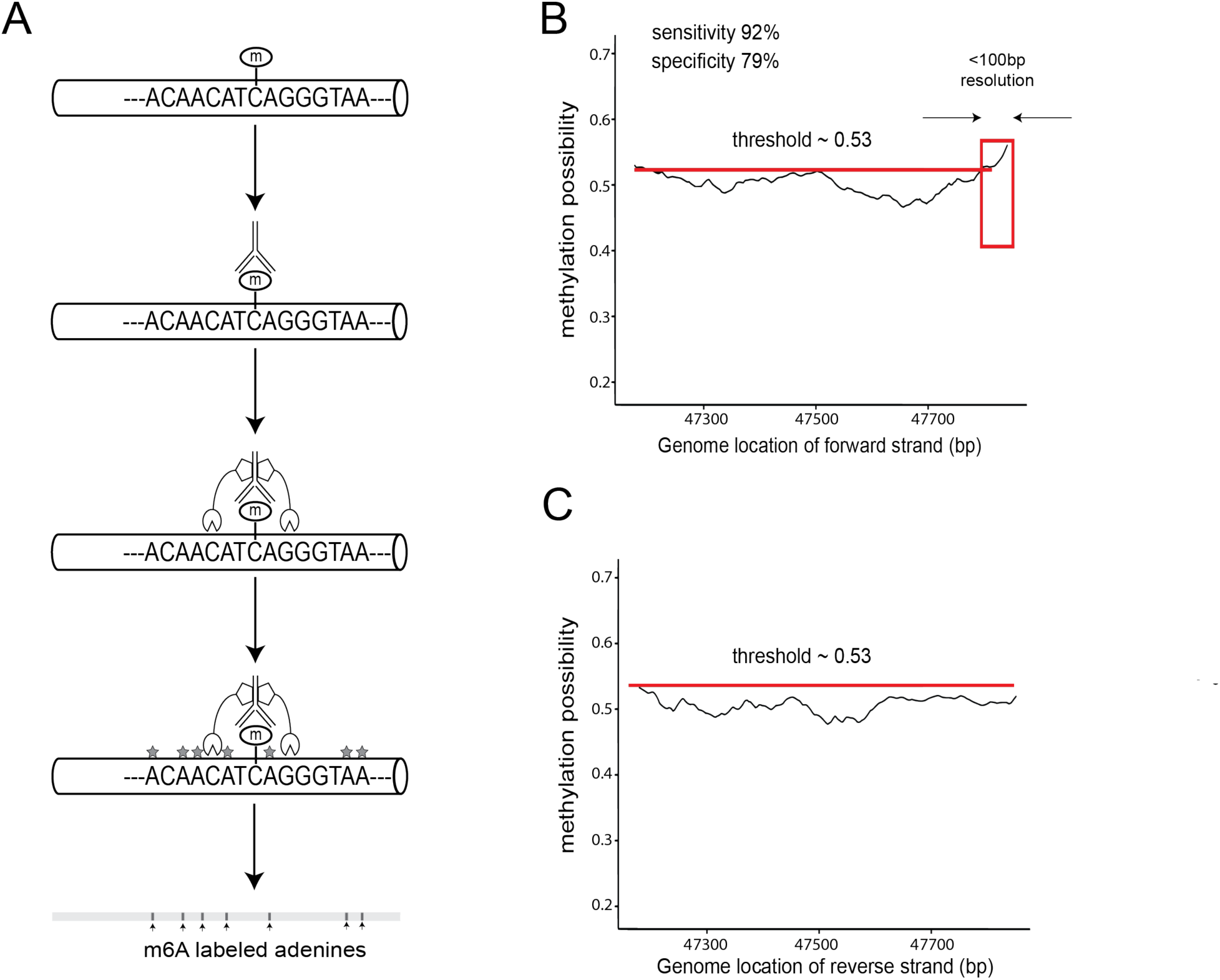
The experimental validation of the m6A calling in BIND&MODIFY *in vitro*. (A) Schematic outline of *in vitro* validation experiment. A fragment of 700bp lambda DNA was amplified by PCR, and 5mC was introduced at 5’ end of forward strand only. The 5mC labeled DNA was bound by 5mC antibody and was subsequently treated by BIND&MODIFY method. (B) The m^6^A possibility (Megalodon calling possibility) of forward strand. When the methylation cut-off was set at 0.53 (red line, sensitivity=0.92, specificity=0.79), high methylation possibility region was observed in 3’ end at resolution of about 100bp, which was originally marked by 5mC at 3’ end. (C) The methylation possibility of reverse strand. No high methylation possibility region was observed above 0.53 cut-off.

In Figure 2B, the m^6^A possibility were plotted along with their genomic location as an indicator of bound pA-M.EcoGII. The regions with high methylation possibility (>0.53, Supplemental Figure 2) corresponded to the neighboring areas of the bound pA-M.EcoGII, where the tethered M.EcoGII could methylate the genome sites within a close distance (Figure 2B). The resolution for detecting pA-M.EcoGII binding sites was around 100bp (Figure 2B), comparable to the conventional ChIP-seq peak size distribution of 100-300bp (Supplemental Figure 3).

The secondary objective of the present study was to investigate the strand specificity of the pA-M.EcoGII modifications. We plotted the methylation possibility, as the indicator of the pA-M.EcoGII location, on both positive and negative strands (Figure 2C). We observed that the pA-M.EcoGII binding signal was located exclusively on the negative strand rather than the positive strand. This suggested that the tethered M.EcoGII was in close proximity to the negative strand of DNA and thus preferentially modified that strand. These features make the BIND&MODIFY method optimal for long-range histone modification studies on the single molecular level and histone modification in a strand-specific manner.

### BIND&MODIFY reveals the comparable strand-specific view of the epigenomic regulator on genomic DNA

ChIP-seq, CUT&RUN, and CUT&TAG detect protein–DNA binding events and chemical modifications of histone proteins ^38–40^. Several histone trimethylation states, such as H3K4me3, are well-studied and have been linked to active gene transcription, whereas H3K27me3’s relationship with transcription status is more ambiguous ^41, 42^. Therefore, we chose to explore the transcription status of H3K27me3 status in the breast cancer cell line MCF-7, using ChIP-seq and BIND&MODIFY. The average read length of Nanopore was around 2kb (Supplemental Figure 4), and the correlation of the experimental replicates was 0.90 (Supplemental Figure 5).

To validate the H3K27me3 signal as determined by BIND&MODIFY, we first evaluated the consistency between that obtained from BIND&MODIFY and the conventional methods. The signals ascertained by both the conventional method (ChIP-seq) and the proposed BIND&MODIFY method provided the similar H3K27me3 position signals in various genome scales (Figure 3A, Supplemental Figure 6). Over 95% of the high confidence position signals gathered using ChIP-seq were also observed using BIND&MODIFY (Supplemental Figure 7). By analyzing the regional signal strength, we found that the BIND&MODIFY signal strength for m^6^A counts was strongly correlated with ChIP-seq peak signal intensity (Figure 3B). Taken together, these BIND&MODIFY results exhibited a range of values comparable with conventional ChIP-seq.

**Figure 3.**
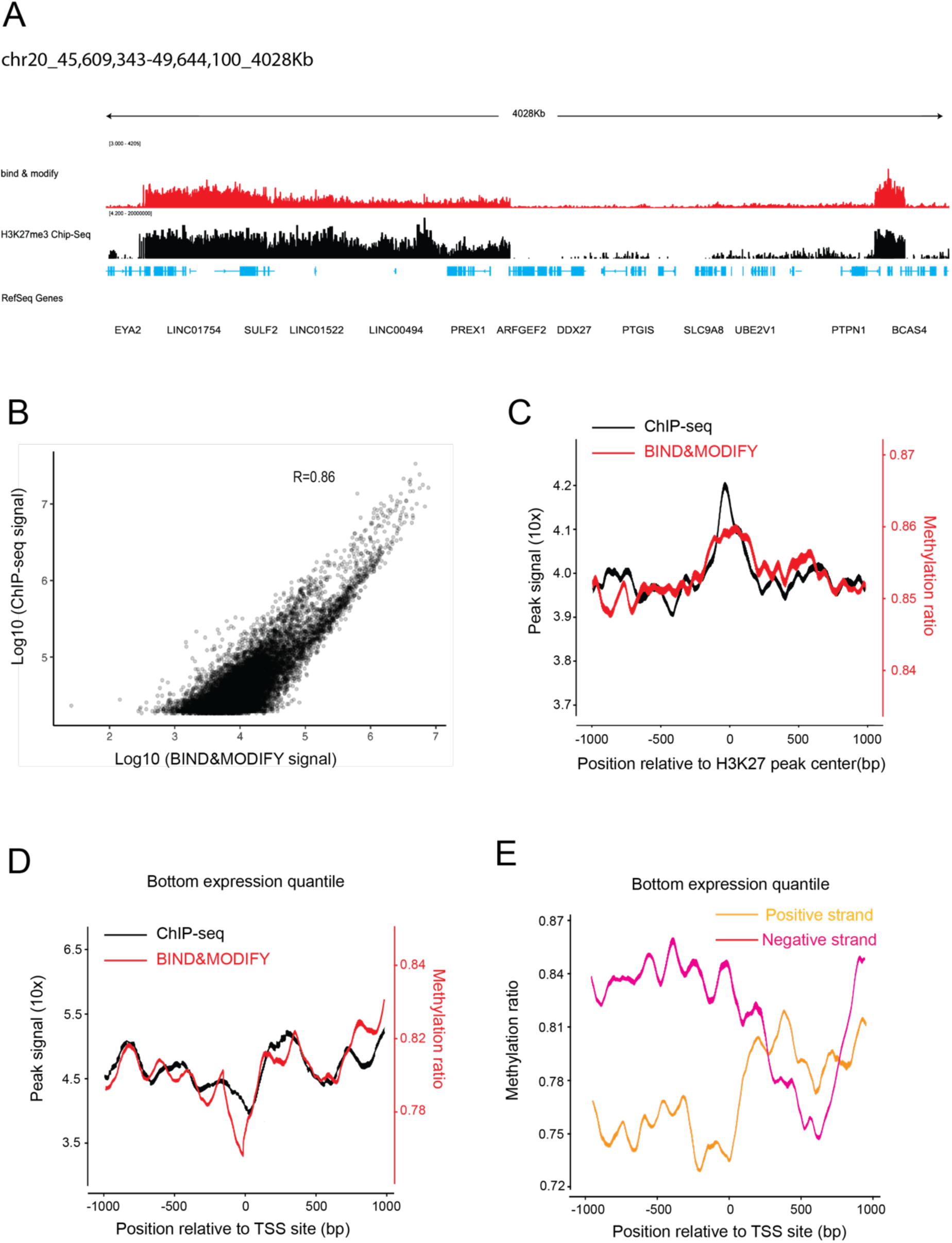
The consistency of H3K27me3 pattern between ChIP-seq and BIND&MODIFY in *vivo*. (A) The H3K27me3 signal, by BIND&MODIFY and ChIP-seq, in genome scale view. (B) The scatter plot of BIND&MODIFY signal and ChIP-seq signal in peak regions. The peak regions are identified H3K27me3 peaks in ChIP-seq. The BIND&MODIFY signal means the m6A counts and the ChIP-seq signal means the read counts. (C) The H3K27me3 pattern of BIND&MODIFY methylation ratio and ChIP-seq signal was plotted by the H3K27me3 peak centered plot (H3K27me3 peaks identified in ChIP-seq) for all the genes on Chr20. The plot covered the upstream/downstream 1000bp from H3K27me3 peak center. (D) The H3K27me3 pattern of BIND&MODIFY methylation ratio and ChIP-seq signal was plotted by the TSS centered plot for the low expression genes. The BIND&MODIFY line was the fitting curve based on the positive strand and negative strand contribution on each region. The low expression genes are the genes in the bottom expression quantile. (E) The H3K27me3 strand-specific view by the TSS centered plot for the low expression genes.

Some studies have observed an inverse relationship between H3K27me3 density in transcription start sites and gene expression ^41, 42^. To profile the enrichment of H3K27me3 within genomic regions, enrichment data were visualized using ChIP-seq H3K27me3 peak centered signal plots. Overall, the BIND&MODIFY results are consistent with those of ChIP-seq (Figure 3C). In the TSSs of low expression genes (bottom expression quantile), H3K27me3 was prominently depleted around the TSS with a distinctive enrichment directly downstream and upstream of the TSSs (Figure 3D). The bimodal pattern is relatively consistent in both ChIP-seq and BIND&MODIFY results. Good agreement was also found when comparing results from this work against published data ^43 44^. Further analysis was performed to determine whether the histone trimethylation exhibited strand specificity using the BIND&MODIFY method. Subsequently, we found that H3K27me3 was prominently enriched downstream of TSS in the positive strand and upstream of TSS in the negative strand, contributing to form the bimodal pattern around TSSs (Figure 3E). An example of the gene is illustrated in Supplemental Figure 8. In contrast, we did not observe the significant strand-specific pattern around the TSSs of high expression genes (Supplemental Figure 8).

As a critical regulator of genome organization, the CCCTC-binding factor (CTCF) has been characterized as a DNA-binding protein with essential functions in maintaining the topological structure of chromatin and inducing gene expression ^45^ (Supplemental Figure 9A). In CUT&TAG data, we identified the strong CTCF binding sites 100bp upstream of the TSSs of actively expressed genes (Supplemental Figure 9B). In contrast, the CTCF binding signal around the TSSs of inactive genes was distributed evenly (Supplemental Figure 9C). Compared with CUT&TAG, BIND&MODIFY presented a similar CTCF signal pattern around TSS in active and inactive genes. Further analysis was done to see whether the antibody-bound CTCF domains maintained selective proximity to either one strand (Supplemental Figure 9D). We found that the CTCF signal on the negative strand was sharper and more potent than on the positive strand and even more consistent with the CUT&TAG signal pattern (Supplemental Figure 9E), suggesting the antibody-bound domain was in close proximity to the negative strand. Based on protein crystal structure analysis ^46^, the antibody-bound domain is the C terminal of the CTCF and spatially closer to the negative strand, further supporting this hypothesis (Supplemental Figure 10). These results have demonstrated that the BIND&MODIFY method has exhibited that it is capable of mapping the histone modifications in a manner comparable with conventional methods, in addition to highlighting the challenges associated with a strand-specific view of epigenomic regulators on genome DNA

### Long-range sequencing resolves the retrotransposon regions with high resolution

The retrotransposon composed of repeated sequences can be integrated elsewhere in a genome and has performs various critical biological functions. The three major retrotransposon orders are long terminal repeat (LTR) retrotransposons, long interspersed elements (LINEs), and short interspersed elements (SINEs). Due to the repeated sequence of retrotransposon, the short-read sequencing data exhibits 40∼60% multiple mapping rate in these complex regions (Supplemental Figure 11). The multiple mapping in retrotransposons would cause signal noise or signal loss in the ChIP-seq (Figure 4A). Compared with ChIP-seq, the BIND&MODIFY significantly improves the unique mapping rate (Supplemental Figure 11), by taking advantage of long-range sequencing, which generates reads that may span the entire length of full transposon and avoid multiple mapping issues. Considering these advantages, the long-range sequencing was also used to identify the novel transposon deletion and insertions^47^. There have been minimal studies regarding the H3K27me3 on retrotransposons ^48^. We visualized the H3K27me3 status on LTR, SINE, and LINE using retrotransposon-centered plots, including an upstream/downstream 300bp of similar size retrotransposons (Supplemental Figure 12). In the ChIP-seq, we observed the very high background noise lacking any peak signal on LTR regions (Figure 4B). In contrast, the BIND&MODIFY displayed the H3K27me3 binding signal on the LTR region, providing a 10X better signal resolution compared to ChIP-seq (Figure 4B). While analyzing another type of retrotransposon, SINE, ChIP-Seq lost the H3K27me3 signal in SINEs. However, BIND&MODIFY displayed two clear peak signals in the SINEs (Figure 4C-D). In addition, these results are consistent well with existing studies regarding H3K27me3 activity on SINEs ^48^.

**Figure 4.**
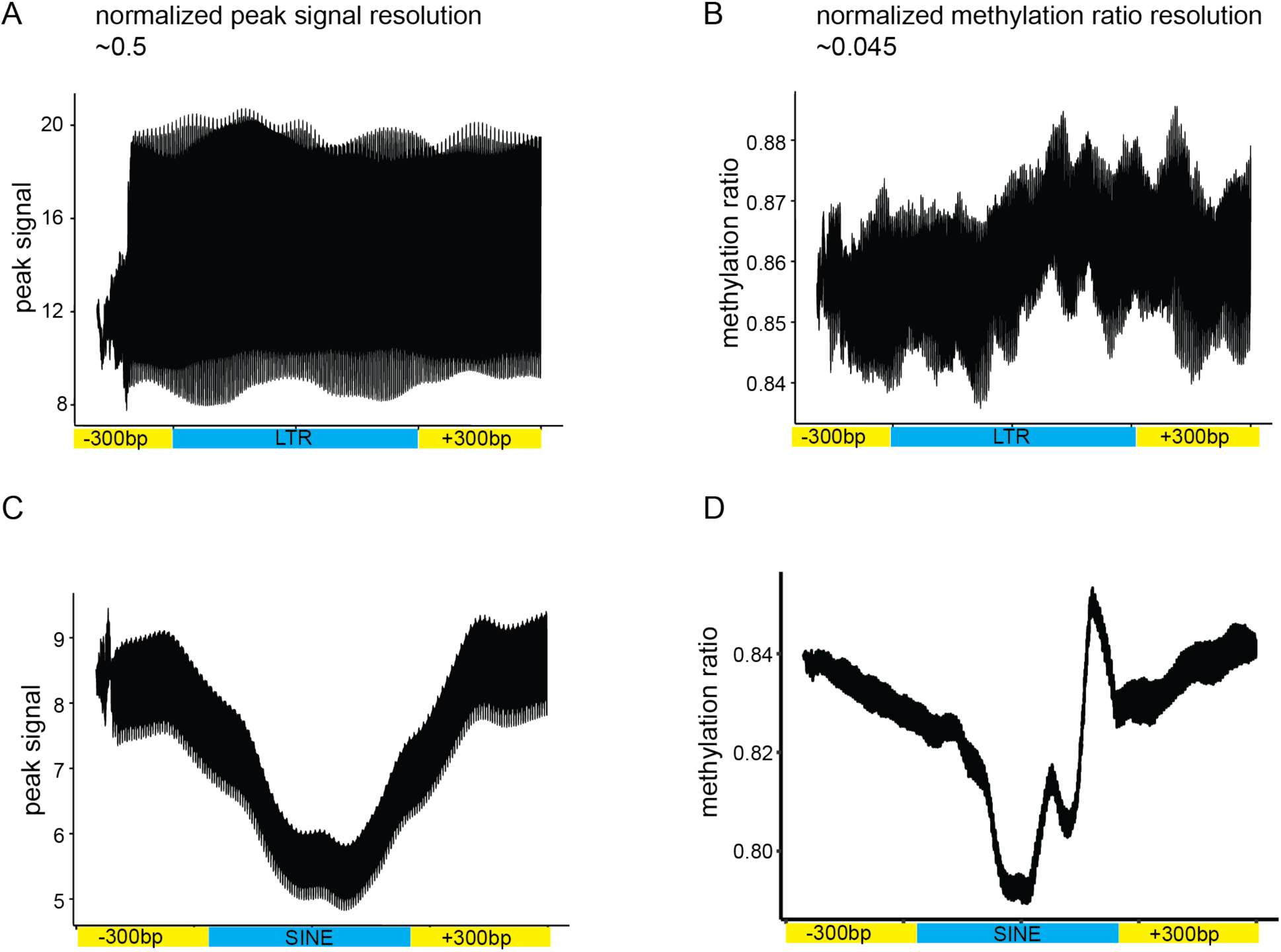
The BIND&MODIFY resolved H3K27me3 in the transposon areas with higher resolution. (A-B) The LTRs with size 350∼450bp were selected and centered. The moving average H3K27me3 signals on the upstream/downstream 300bp of these LTRs, including LTRs, were plotted with corresponding genomic sites. The y-axis (A) indicated the normalized read counts of ChIP-seq. The y-axis (B) indicated the normalized methylation ratio of m6A in BIND&MODIFY with nanopore sequencing. Normalized peak signal resolution is defined by ratio of mean peak signal width (maximum-minimum) and mean maximum peak signal, which is about 0.5 in ChIP-seq and 0.045 in BIND&MODIFY. (C-D) The SINEs with size 250∼350bp were selected and centered. The moving average H3K27me3 signal on the upstream/downstream 300bp of these SINEs, including SINEs, were plotted with corresponding genomic sites. The y-axis (C) indicated the normalized read counts of short-reads ChIP-seq. The y-axis (D) indicated the normalized methylation ratio of m6A in BIND&MODIFY with nanopore sequencing.

The short interspersed nuclear element (SINE) B1 has insulator activity mediated by the binding of specific transcription factors along with the insulator-associated protein CTCF ^49^. A genome-wide analysis of CTCF binding sites in the human and mouse genomes discovered that many CTCF binding sites are derived from TE sequences ^50^. Concordant with previous reports, the BIND&MODIFY exhibited a clear CTCF signal in the SINEs, which was absent in the ChIP-Seq data (Supplemental Figure 13). In summary, BIND&MODIFY is capable of taking advantage of the long-range sequencing to precisely map histone modifications and protein-DNA interactions in the challenging complex genome regions.

### The histone modification status on single-molecular resolution

The conventional CUT&TAG and ChIP-seq methods are based on statically calling the peak of the enriched read in a specific region ^51^. Recent single-molecule and single-cell measurements of histone accessibility suggest that using ATAC-seq to evaluate cell populations represent an ensemble average of distinct nucleosome states ^52^. An essential attribute of the BIND&MODIFY technique, is that it measures the histone modification in single-molecular resolution by taking advantage of the slight variance (Supplemental Figure 14), thereby increasing the cumulative possibility of segments (Supplemental Figure 15).

We then hypothesized whether BIND&MODIFY could reveal the different H3K27me3 statuses by investigating the chr20:52223000-52225500 loci. The chr20:52223000-52225500 loci have modulated the lncRNA LOC105372672 and Zinc finger protein 217 (ZNF217) and has been demonstrated as a prognostic biomarker and therapeutic target during breast cancer progression ^53, 54^. The conventional ChIP-Seq enriched the H3K27me3 bound motif by antibody-guided amplification, which could be overrepresented (Figure 5A).

**Figure 5.**
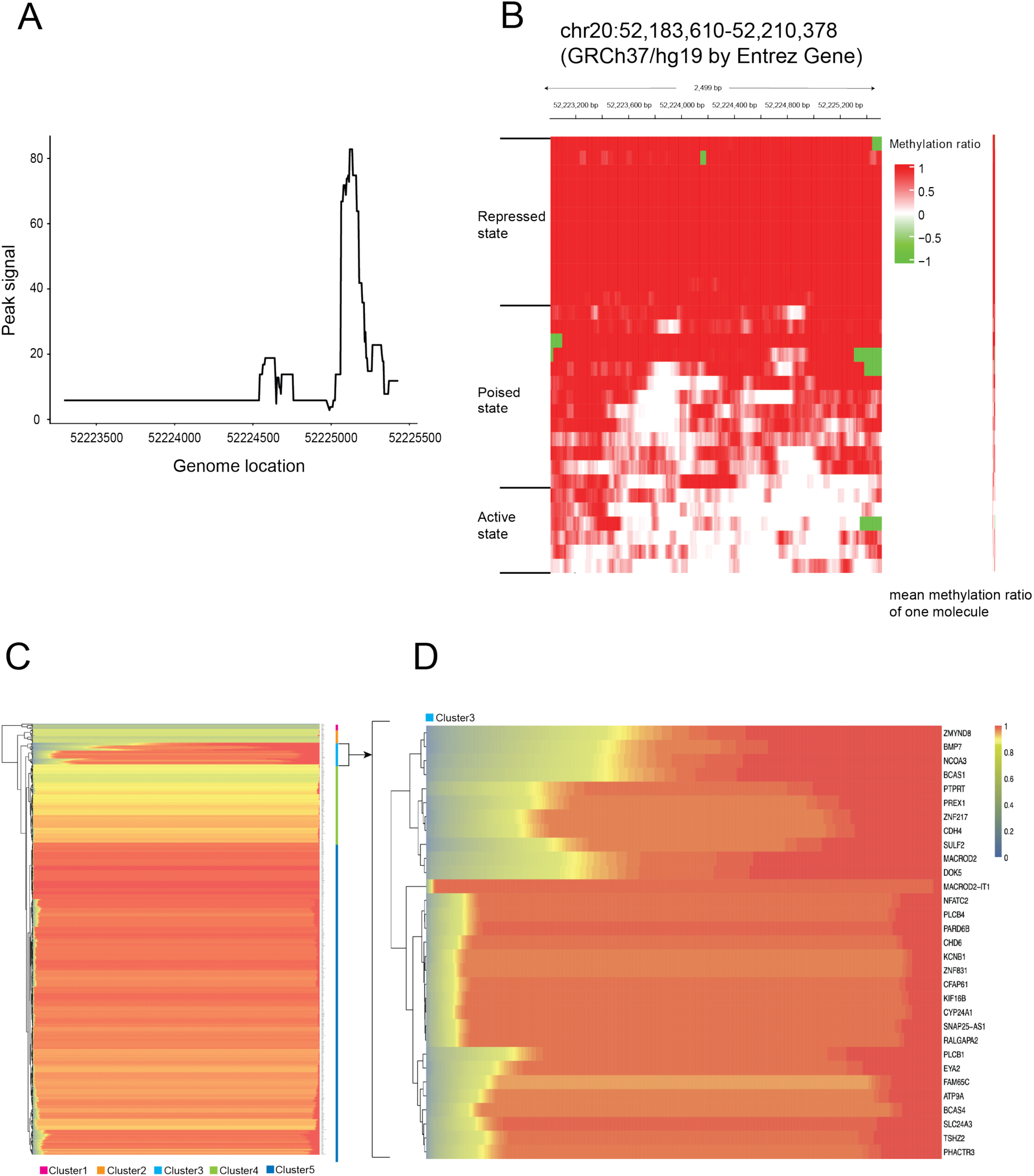
The BIND&MODIFY showed the heterogeneity of H3K27me3 regulation. (A) The peak signal of H3K27me3 in ChIP-seq (Chr20:52,183,610-52,210,378). (B) The single molecular resolution of the positive strand Chr20:52,183,610-52,210,378 visualize each molecular methylation statues of H3K27me3. The rows in heatmap represented the different DNA molecules. The color indicated the methylation ratio, which represented the H3K27me3 signal. The DNA molecules could be classified into three states: repressed state, poised state, and active state based on their mean methylation ratio. The right panel is the condensed illustration of individual DNA molecules mean methylation ratio. Each block on the vertical line (molecule heterogeneity line) represented the mean methylation of each DNA molecule. (C) H3K27me3 heterogeneity pattern of genes in chr20. Mean H3K27me3 methylation ratio of individual DNA molecule was plotted for all the genes on DNA region (Chr20:52,183,610-52,210,378). Each pixel on each row corresponds to mean methylation ratio of each individual DNA molecule, and each row corresponds to each gene. The methylation ratio was ranked from low to high (left to right). (D) The right heatmap showed the magnified cluster 3 in the left heatmap. Details of H3K27me3 heterogeneity calculation methods can be found in supplementary figure 17.

The baseline signal of this region was 6, which was considered to be the background noise in the experiments (Figure 5A). In contrast, the BIND&MODIFY presents a strikingly comprehensive picture of the H3K27me3 in this area. Remarkably, super-resolution of BIND&MODIFY uncovered remarkable three different epigenetic states (Figure 5B): a repressed state in which most histones were methylated, inhibiting the gene expression contained within the genome region; a poised state in which half of the histones were methylated, which permit the transition from the inactive state to the active state; and an active state largely devoid of histone trimethylation. Some of the H3K27me3 could only be observed in the subpopulation of the genome fibers, suggesting the highly heterogenous histone methylation in cancer cells. The signal peaks (chr20: 52225000-52225500) in ChIP-seq sum up the H3K27me3 loci in two different states. The subpopulation of inactive fully methylated chromatin leads to the baseline signal in ChIP-seq, which was thought to be the background noise in the experiment. Our findings, which support three epigenetic states, align with the results reported by other publications in literature ^55–58^. Many genome regions in cancer cells harbor a distinctive histone modification signature that combines the activating histone H3 Lys 4 trimethylation (H3K4me3) mark and the repressive H3K27me3 mark. The poised states with these bivalent domains, which are considered to be linked to poise the expression of developmental genes, permitting timely activation (activate form) while maintaining repression (repressed state) in the absence of differentiation signals ^55^. In contrast to the heterogeneous behaviors seen above, the gene desert areas without gene transcription activity processed only one repressed state wherein most histones were trimethylated (Supplemental Figure 16). The previous bivalent model was established by finding the overlapped H3K4me3 and H3K27m3 average peaks in NGS but not acknowledge their poised status. These findings reinforce the general belief that bivalent histone modification regulates gene expression.

However, the single molecular statuses of each gene are challenging to be quantified. Therefore, we could not obtain a big-picture view of global genes, summarizing a more comprehensive perspective with specific examples and details. Consequently, we used the mean methylation ratio of one molecule to present the methylation status of this molecule (Supplemental Figure 17), and the heterogeneity of the genes could be visualized by a series of mean molecular methylations (Figure 5B, gradient line on the right panel). We summarized the H3K27me3 heterogeneity of genes in Chr20 (Figure 5C). 95% of genes display a homogenous H3K27me3 regulation status with minimal heterogeneity among molecules. Only one cluster with 31 genes (Figure 5C, cluster 3) adequately demonstrated the very high heterogeneity of H3K27me3 regulation, illustrating its active, repressed, and poised state. These genes are under typical bivalent mode regulation of H3K27me3. Using gene ontology analysis, we found that these genes were enriched in the breast cancer-related pathways, for example, the VEGF singling pathway^59^, B cell receptor signaling pathway^60^, PD-L1 checkpoint pathway^61^, etc. (Supplemental Figure 18). The bivalent mode regulation in these pathways might elucidate the mechanisms involved in tumor heterogeneity, evolution, drug resistance, immune evasion, and the cause of metastasis.

### The BIND&MODIFY reveals the long-distance correlation of regulators

In the transition from a chromatin repressed state to an activated state, the H3K27me3 changed its rhythmics. The region (chr20: 52223500-52224200) was first devoid of H3K27me3 and then adjacent areas concurrently removed any histone trimethylation during the next step, suggesting the possibility of a synergic regulatory switch at this location. To quantify distance (anti)correlation between chromatin trimethylation states, we developed a modified correlation coefficient (CC) metric for assessing the degree of trimethylation correlation between genomic regions. Average CC profiles centered on statically positioned nucleosomes revealed a detectable correlation between nucleosome positions up to 100bp (Supplemental Figure 19). These observations are consistent with the assumption that nucleosomes impose restrictions on one another, resulting in a short-range correlation between nucleosome footprints that dephases over longer ranges ^10^. Furthermore, CC analysis of DNA confirmed the long-range positive correlation between the promoter region and this upstream, downstream element (Figure 6A, Supplemental Figure 20).

**Figure 6.**
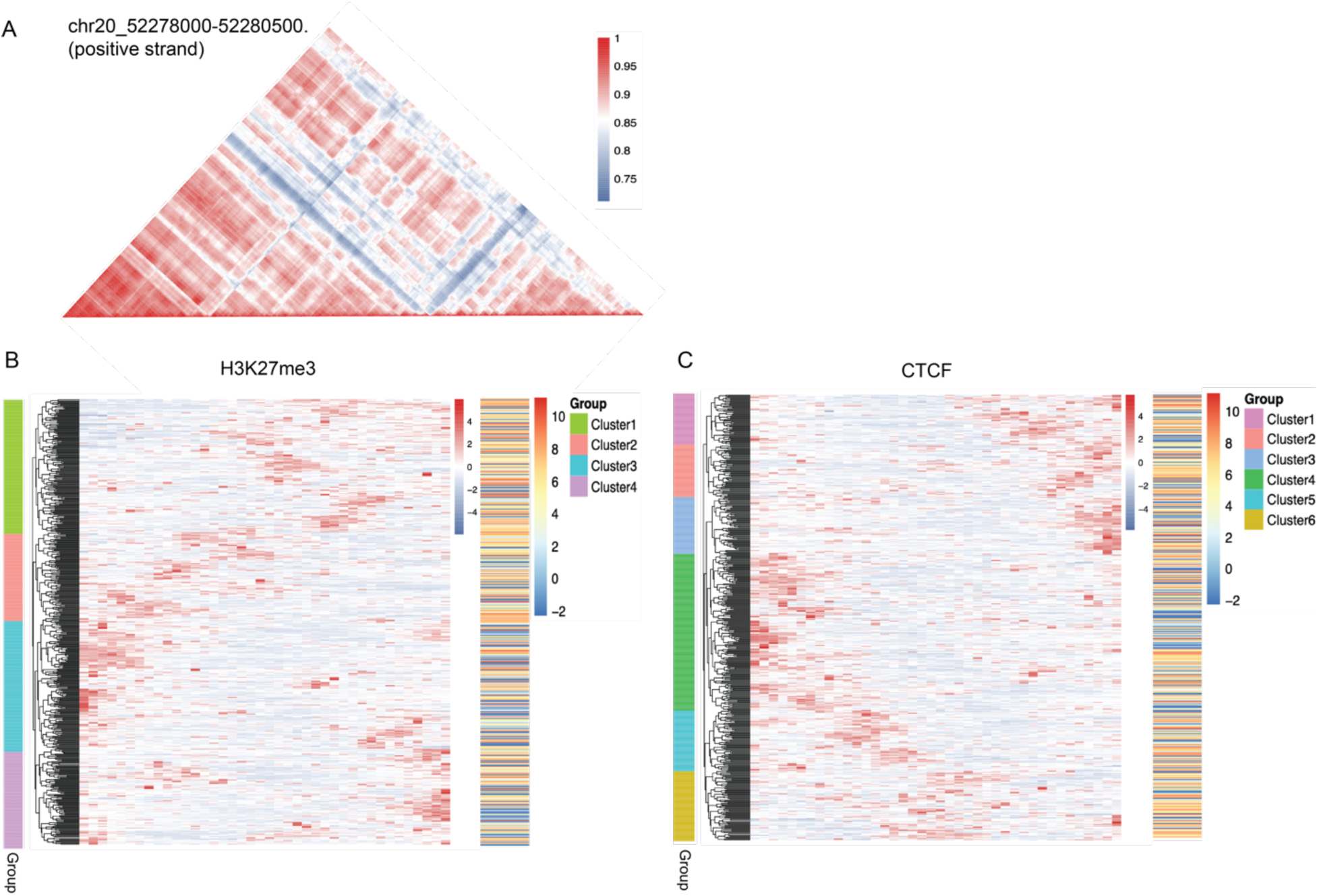
The distance correlation of the epigenetic regulation. (A) The distance correlation of the H3K27me3 in chr20: 52278000∼52280500 (positive strand). The color indicated the correlation coefficient (CC) metric among genome regions. The higher CC metric means the stronger correlation in distance. (B) The global view of the H3K27me3 distance effect (DE) index for all the genetic transcription regions (upstream 2kb of transcription start site) in chr20. The DE index is sum of the CC*distance, representing this location impact on other sites. The red color means the stronger impact on distal genomic sites. The left color bar indicated the clusters of these transcription regions. The right color bar indicated the corresponding gene expression. (C) The global view of the CTCF distance effect for all the genetic transcription in chr20. Details of correlation coefficient (CC) metric and distance effect (DE) index calculation method can be found in supplementary figure 21.

However, the CC index is only suitable for describing the distance correlation of H3K27me3 for single genes. To further explore the biological significance of distance correlation, we developed the new DE index (Distance Effect index) to quantify the distal effects of each genomic locus (Supplemental Figure 21). The higher DE index means that these target genomic loci have a stronger distal correlation, suggesting that the regulatory mechanisms present on these sites could impact a larger area. Therefore, the genetic region of interest could be presented as a series of DE indices on the corresponding loci. We plotted the DE index of all the transcription regions (2kb upstream of TSS) on chr20 (Figure 6B). There were several DE patterns consistent with H3K27me3 in the transcription region. Some transcription initiation sites had a powerful DE index signal, which suggested the rhythmic change in the more extensive area and its substantial regulator impact on the distal genomic region (Figure 6B). Consistent with studies that have associated H3K27me3 with transcriptional repression ^62^, a strong DE index signal for H3K27me3 was related to lower gene expression (Figure 6B). Based on gene ontology analysis, the dysregulation of these genes was associated with cancer pathways, such as those for ubiquitin dysregulation, Apelin signaling, proteasome, Oxytocin signaling, etc (Supplemental Figure 22).

We further plotted the distance correlation of transcription regions with CTCF regulators. The the areas where the transcription initiation sites have strong DE index signal (Figure 6C, Cluster 3) indicate that the CTCF on those sites may synergically affect the distal genomic sites. Compared with the gene expression profile, some gene expression is strongly enhanced. By gene ontology analysis, these enhanced distance regulations were also related to the cancer-promoted pathways such as those for EGFR tyrosine kinase inhibitor resistance, GABAergic synapse, Hippo signaling pathway, etc. (Supplemental Figure 23). In addition, the observed enhanced distance regulation may have elucidated the super-enhancer/suppressor effects, which do affect the gene region up to 8kb ^1^.

Overall, the BIND&MODIFY was able to demonstrate the highly heterogenous status of histone modification in the cancer cells and the long-range interaction of these histone modifications.

## Discussion

In literature, ChIP-seq and CUT&TAG have been essential epigenetic study tools for many years, for example, histone modification, transcription factor binding, and much more. However, these methods suffer from the limitations that the detected signals are represented using common immunoprecipitation enriched DNA fragments in common without considering the heterogeneity in DNA molecules. The short-read sequences also prevent the possibility of studying the long-range interactions in multi-omics epigenome studies. We designed a new method, “BIND&MODIFY,” by using the recombinant pA-M.EcoGII methyltransferase to non-fragmented labeling the local DNA regions. Through this method, we were able to simultaneously profile multi epigenomic information on a truly unbiased genome-wide scale, measure the underlying distribution of the histone modifications based on single molecular resolution, and identified loci exhibiting significant correlation.

BIND&MODIFY generated a similar general signal trend for the H3K27me3 loci to the widely used ChIP-seq. Despite the significant commonalities between the results of the two methods, relatively small signal variations were observed in terms of the variable peak signal strength. We postulate that these were likely caused by the different fundamental principles between BIND&MODIFY and conventional ChIP-seq. The BIND&MODIFY enables unbiased profiling of the DNA molecule without immunoprecipitation enrichment, thereby reflecting the true nature of the epigenome. Compared with that method, ChIP-seq only amplifies the common loci signals among most DNA molecules by immunoprecipitation enrichment and neglects the loci signals of in individual DNA molecules.

Moreover, the different measuring units to present in signal strength also led to the difference. One drawback of the BIND&MODIFY is that it sequences the non-signal regions, which is unavoidable, thereby requiring a higher sequencing depth. Fortunately, the nanopore throughput is increasing rapidly, while selective enrichment methods are also becoming increasingly available. Augmenting the read length are also helpful, especially for analyzing the distance correlation of distal regulatory elements.

We also compared our method with CUT&TAG. The significant difference is that the transposon utilized in CUT&TAG is not reusable once inserted into the DNAs. In contrast, the methyltransferase used in BIND&MODIFY could methylate the multiple regional adenosines, enhancing the signal strength. CUT&TAG also requires a secondary antibody to tether more transposons to the location and increase the chance of DNA insertion. In BIND&MODIFY method, we found that the secondary antibody is not necessary. Others may suggest that the reusable methyltransferase may increase the false-positive signals in non-target regions. However, during *in vitro* evaluation, we found the only scattered signals in the non-targeted area, but no clustered signals, which were consistent with identifiable loci signal (Figure 2). In addition, during *in vivo* evaluation, the genomic DNA was not movable by fixation, and the DNA methyltransferase can only activate locally (Figure 3). Therefore, any false-positive signal of *in vitro* and *in vivo* can be safely neglected based on these studies.

Base-calling is another area of future improvement. We also used the Pacbio sequencing data (specificity 0.99) to train our m^6^A calling algorithm with nanopore sequencing. Both sensitivity and specificity with regard to nanopore m^6^A detection were satisfactory. We are also trying to call the native m^5^C and artificially labeled m^6^A simultaneously and analyze the relationship between histone modification and DNA methylation. To our surprise, the artificially labeled m^6^A did not significantly affect the m^5^C detection efficiency significantly, and the m^5^C correlation using bisulfite sequencing was 0.8 (Supplemental Figure 24). The relationship between 5mC and histone modification in single molecules should be determined in much deeper sequencing depth. Alternatively, we could also use the Pacbio, which reads one nucleotide at a time without the neighboring nucleotide signal interference. Overall, the simultaneous detection of 5mC and its regulator position in the single-molecule level is applicable with BIND&MODIFY.

Possible endogenous methylation in mammalian genomes also represents the potential confounding factor in our analysis. By examining the data of IgG control genomic DNA, we found that the quantity of m^6^A in original DNA is thousands of times fewer than in the treated samples, suggesting the minimal effect of the endogenous methylation. To determine the thresholds for the possibility of real m^6^A and background noise, the IgG control sample data was used to determine and filter out the background noise signal distribution is below 0.53, which could significantly improve the m^6^A calling specificity and sensitivity.

However, there are also some species where m^6^A occurs endogenously and strongly correlated with the binding motif of the histone modification. Modifications targets such as m^5^C, m^4^C, cytidine deamination, or 5-glucomethyaltion are among the potential future alternatives in such cases. Also, some artificial SAM could be used to introduce the biotin to the specific sites may strongly avoid the endogenously confounding signals. Finally, we believe that the integration of BIND&MODIFY into a single-molecule multi-omics assay represents a fruitful direction that will benefit future research. With the method, we are able to simultaneously label the two or more proteins located in the single DNA molecule with the different modification labels. Then the protein interaction distance on the genome would be measurable in this method, providing a potential breakthrough in the protein-protein interaction in single-molecule genome scales. In principle, similar approaches may also apply to the individual RNA molecules to study the RNA binding proteins. We expect this technology to contribute to the essential new class of tools that will improve the simultaneous study of multi-epigenomics.

## Methods

### Cell culture and antibodies

Human mammary gland carcinoma cell line MCF-7 were obtained from ATCC. MCF-7 were grown in DMEM (Gibco,11995065) supplemented with 10% FBS (Gibco,10099141), 0.01mg/ml insulin (HY-P1156, MedChemExpress), and 1% penicillin-streptomysin (Gibco, 15140122). Cell line was regularly checked for mycoplasma infection (Yeasen, 40612ES25). We used the following antibodies: H3K27me3 (Cell Signaling Technology, 9733), Guinea Pig anti-Rabbit IgG (Heavy & Light Chain) antibody, Anti-CTCF Antibody (sigma,07-729-25UL), Protein A Antibody, pAb, Chicken (A00729-40, GenScript).

### Recombinant protein preparation of protein A-M.EcoGII

pTXB1 vector (NEB, N6707S) was used as the protein expression backbone. Downstream of the lac operator, a ribosome binding site and three FLAG epitope tags were introduced, followed by two IgG-binding domains of staphylococcal protein A encoding sequence, which was synthesized based on the previous work (Addgene, 124601). The M.EcoGII encoding sequence (Addgene, 122082) was also synthesized based on its original discovery. The amino acid linker sequence between the C-terminus of protein A and N-terminus of M.EcoGII is DDDKEF. The sequenced plasmid was transformed into C3013 competent cells (NEB) following the manufacturer’s protocol. Each colony tested was inoculated into a 1 mL LB medium, and growth was continued at 37 °C for 2 h. That culture was used to start a 50 mL culture in 100 μg/mL carbenicillin containing LB medium and incubated on a shaker until the cell density reached an A600 of 0.6, whereupon it was chilled on ice for 30 min. Fresh IPTG was added to 0.25 mM to induce expression. Then the culture was incubated at 27 °C on a shaker for 16 h. The culture was then collected by centrifugation at 10,000 rpm, 4 °C for 30 min, the supernatant was discarded. The bacterial pellet was frozen in a dry ice-ethanol bath and stored at −70 °C. The frozen pellet was resuspended in 20 mL chilled HEGX Buffer (20 mM HEPES-KOH at pH 7.2, 0.8 M NaCl, 1 mM EDTA, 10% glycerol, 0.2% Triton X-100) including 1× Roche Complete EDTA-free protease inhibitor tablets and lysed using a high-pressure cell disrupter (JNBIO, China). Cell debris was removed by centrifugation at 10000 rpm for 30 min at 4 °C, and the supernatant was loaded onto a column equipped with chitin slurry resin (NEB, S6651S), then incubated the column on a rotator at 4 °C overnight. The unbound soluble fraction was drained, and the columns were rinsed with 20 mL HEGX containing Roche Complete EDTA-free protease inhibitor tablets. The chitin slurry was transferred to a 15 mL conical tube and resuspended in Elution buffer (10 mL HEGX with 100 mM DTT). The tube was placed on the rotator at 4 °C for ∼48 h. The eluate was collected and concentrated using an Amicon Ultra-4 Centrifugal Filter Units 10 K (Millipore, UFC801096), and sterile glycerol was added to make a final 50% glycerol stock of the purified protein. The fusion protein has stored the protein at -80°C.

### Size characterization of recombinant protein pA-M.EcoGII

The size of purified pA-M.EcoGII recombinant protein was characterized by C Coomassie blue staining by resolving the protein in 7.5% SDS-PAGE gel.

### ELISA of recombinant protein pA-M.EcoGII

To verify the binding efficiency of pA-M.EcoGII, we performed an ELISA assay in vitro. The purified recombinant protein or commercial M.EcoGII (NEB, M0603S) was diluted with coating buffer (0.05 M NaHCO_3_ buffer, pH 9.2). The 96 well high binding plate (Greiner Bio-one, 655061) was coated with 100 μL pA-M.EcoGII or commercial M.EcoGII (NEB,M0603S) (1:120, 1:480 of stock 10mg/ml), negative control, commercial ProteinA (ThermoFisher, 21181) (1:120, 1:480 of stock 10mg/ml) in coating buffer per well for 4 h at room temperature. Further, 200 μL SuperBlock™ (TBS.) Blocking Buffer – Blotting (Invitrogen,37537) was added to each well and incubated for 2 h at room temperature. Then each well was washed 5X with 380 μL washing buffer (0.14 M NaCl; 0.01 M PO_4_ ;0.05% Tween 20; pH 7.4). After that 95 μL Secondary Antibody (1:10000 in PBSB) was added to each well and incubated for 1 h at room temperature. The plate was washed 4X with 380 μL washing buffer (0.14 M NaCl; 0.01 M PO_4_ ;0.05% Tween 20; pH 7.4). 90 μL TMB substrate solution was added and incubated for 15min at room temperature in the dark. Finally, 90 μL stop buffer (1.8N H_2_SO_4_) was added to stop the color development and read immediately at 450nm (yellow color) using FLUOstar® Omega Plate Reader by BMG LABTECH.

### *In vitro* methtransferase activity of pA-M.EcoGII recombinant protein

To access the in vitro methylation efficiency of pA-M.EcoGII recombinant protein, m^6^A methylation-sensitive restriction enzyme DpnI was used to probe the adenine methylation at GATC motif of 7 kb unmethylated dsDNA. The 7 kb dsDNA substrate was PCR amplified from lambda DNA. For the methyltransferase reactions, each 50 μl reaction volume was assembled on ice and contained 1μg 7kb unmethylated dsDNA, 1X Cutsmart buffer, 640 μM SAM, and 4 μL of pA-M.EcoGII recombinant protein or commercial M. EcoGII, then the mixture was incubated at 37 °C for 1 h. The methylated product was purified using 0.6X Ampure XP (BECKMAN COULTER, A63882). 1 μl of DpnI (NEB) was added to the reaction mixture to further incubate at 37 degrees for 10min. DpnI cutting efficiency was examined by 1% agarose gel electrophoresis.

To assess the specificity of pA-M.EcoGII recombinant protein methylation and its effectiveness at inhibiting restriction endonucleases, we carried out restriction analyses using an unmethylated 7kb linear dsDNA template, which was PCR amplified from pTXB1. One enzyme known to be insensitive to dA methylation (BamHI) and three enzymes (EcoRV, PciI, PvuII) that cleave different base-pair sequences, the activities of which are known to be blocked by adenine methylation. The commercial M.EcoGII and pA-M.EcoGII recombinant protein methyltransferase reactions on the 7kb linear dsDNA were carried out the same as described above. All restriction endonucleases used in this study were purchased from NEB. For the restriction enzyme digest reaction, each 30 μL reaction volume contained 500 ng methylated dsDNA, the appropriate digestion buffer, time, and amount of enzyme following the manufacturer’s protocol of NEB. The enzymes were inactivated at 80 °C for 20 minutes. All sample was loaded to 1% agarose gel for analysis.

### *In vitro* pA-Tn5 transposome preparation

The pA-Tn5 was purchased from Vazyme (Vazyme, S603). To generate the pA-Tn5 adapter transposon, 7 μL of a 50 μM equimolar mixture of pre-annealed Tn5MEDS-A and Tn5MEDS-B oligonucleotides, 40 μL of 7.5 μM pA-Tn5 fusion protein, and 28 μL coupling buffer were mixed. The mixture was incubated for 1 h on a Thermocycler at room temperature and then stored at −20 °C.

### CUT&TAG for bench-top application

Gently resuspend and withdraw enough of the slurry such that there will be 10 μL for each final sample. Place the tube on a magnet stand to clear (30 s to 2 min). Withdraw the liquid, and remove it from the magnet stand. Add 1.5 mL Binding buffer (20 mM HEPES pH 7.5, 10 mM KCl, 1 mM CaCl_2,_ and 1 mM MnCl_2_), mix by inversion or gentle pipetting, remove liquid from the cap and side with a quick pulse on a microcentrifuge. Resuspend in a volume of Binding buffer equal to the volume of bead slurry (10 μL per final sample). 10 μL of activated beads were added per sample and incubated at room temperature for 15 min. Cells were harvested, counted and centrifuged for 3 min at 600×g at room temperature. 500000 cells were washed 2X in 1.5 mL Wash Buffer (20mM HEPES pH 7.5, 150mM NaCl, 0.5mM Spermidine, 1× Protease inhibitor cocktail), after that the cells were resuspended in 1.5 mL Wash Buffer by gentle pipetting in a 2mL tube. The unbound supernatant was removed by placing the tube on the magnet stand to clear and pulling off all of the liquid. The bead-bound cells were resuspended with 50 μL Dig-Wash Buffer (20mM HEPES pH 7.5, 150mM NaCl, 0.5mM Spermidine, 1× Protease inhibitor cocktail, 0.05% Digitonin) containing 2mM EDTA and a 1:100 dilution of the appropriate primary antibody. The primary antibody was incubated on a rotating platform overnight at 4 °C. The primary antibody was removed by placing the tube on the magnet stand to clear and pulling off all of the liquid. The secondary antibody was diluted 1:100 in 100 μL of Dig-Wash buffer, and cells were incubated at room temperature for 1h. Cells were washed using the magnet stand twice for 5 min in 1 mL Dig-Wash buffer to remove unbound antibodies.0.04 μM of pA-tn5 was prepared in 150 μL Dig-Med Buffer (0.05% Digitonin, 20 mM HEPES, pH 7.5, 300 mM NaCl, 0.5 mM Spermidine,1× Protease inhibitor cocktail). After removing the liquid on the magnet stand, 150 μL was added to the cells with gentle vortexing, which was incubated with pA-Tn5 at room temperature for 1 h. Cells were washed twice for 5 min in 1 mL Dig-Med Buffer to remove unbound pA-Tn5 protein. Next, cells were resuspended in 300 μL Tagmentation buffer (10 mM MgCl_2_ in Dig-Med Buffer) and incubated at 37 °C for 1 h. To stop tagmentation, 15 μL EDTA,3 μL 10% SDS, 2.5 μL 20mg/ml proteinase K was added to 300 μL of the sample, which was incubated in a water bath overnight at 55 °C. 320 μL PCI was added to the tube and mixed by full-speed vortexing for 2 s. The upper phase was transferred to a phase-lock tube. 320 μL Chloroform was added to the tube and inverted ∼10x to mix. The tube was Centrifuged for 3 min at room temperature at 16,000 x g. The aqueous layer was transferred to a fresh 1.5 mL tube containing 750 μL 100% ethanol and mixed by pipetting. The tube was Chilled on ice and centrifuged for at least 10 min at 4 °C 16,000 x g. The liquid was removed, and 1 mL 100% ethanol was added to the tube, then centrifuged 1 min at 4 °C 16,000 x g. The liquid was carefully poured off and air dry. When the tube is dry, 25 μL 10 mM Tris-HCl pH 8, 0.1 mM EDTA was added to the tube and vortex on full of dissolving the genomics DNA

### BIND&MODIFY for bench-top application

500,000 cells were used in each BIND&MODIFY assay. Cells were harvested, counted, and centrifuged for 3 min at 600×g at room temperature. Cells were first lightly fixed by adding formaldehyde (ThermoFisher, 28906) to a final concentration of 0.1% in 1.5ml PBS, and incubated at room temperature for 15min. 2.5M Glycine was added to final concentration of 0.125 M to quench the additional formaldehyde. Fixed cells were then washed twice in 1.5 mL Wash Buffer (20mM HEPES pH 7.5, 150mM NaCl, 0.5mM Spermidine, 1× Protease inhibitor cocktail) by gentle pipetting. 10 μL of activated beads were added per sample and incubated at room temperature for 15 min. The unbound supernatant was removed, bead-bound cells were resuspended in 100 μL Dig-Wash Buffer (20 mM HEPES pH 7.5, 150 mM NaCl, 0.5 mM Spermidine, 1× Protease inhibitor cocktail, 0.05% Digitonin) containing 2 mM EDTA and a 1:100 dilution of the appropriate primary antibody. Primary antibody incubation was performed on a rotating platform overnight at 4 °C. The primary antibody was removed by placing the tube on the magnet stand to clear and pulling off all of the liquid. No secondary antibody was used for BIND&MODIFY. Cells were washed twice using the magnet stand for 5 min in 1 mL Dig-Wash buffer. 4 μL of pA-M.EcoGII was prepared in 150 μL Dig-Wash Buffer. After removing the liquid on the magnet stand, 150 μL of pA-M.EcoGII containing Dig-Wash buffer was added to the cells with gentle vortexing, which was incubated at room temperature for 1 h. Cells were washed 2× for 5 min in 1 mL Dig-Wash Buffer and 1x for 5 min in 1mL Low salt buffer (20 mM HEPES pH 7.5, 0.5 mM Spermidine, 1× Protease inhibitor cocktail, 0.05% Digitonin) to remove unbound pA-M.EcoGII protein. Next, cells were resuspended in 300 μL Reaction buffer (20 mM HEPES pH 7.9, 0.5 mM Spermidine, 1× Protease inhibitor cocktail, 0.05% Digitonin, 10 mM 1M Mgcl_2_, 300 mM sucrose) .The reaction was activated by adding 5 μL SAM of 32 mM at 37°C in a thermmixer. Additional 5 μL SAM of 32 mM was added to the tube at 7.5 min and 15 min. The reaction was stopped at 30 minutes by placing the tube on the magnet stand to clear and pulling off all of the liquid. The bead-bound cells was resuspended with 300 μL Digestion Buffer(20 mM HEPES pH 7.5, 300 mM NaCl, 0.5 mM Spermidine, 1× Protease inhibitor cocktail, 0.05% Digitonin, 16.7 mM EDTA, 0.1% SDS, 0.167 mg/mL proteinase K) and incubated in water bath overnight at 55 °C. The method for genomic DNA extraction was the same as CUT&TAG for bench-top application.

### CUT&TAG library preparation

To amplify libraries, 21 μL of CUT&TAG genomic DNA was mixed with 2 μL of a universal i5 and a uniquely barcoded i7 primer, using a different barcode for each sample. A volume of 25 μL KAPA HIFI ready mix (KAPA, KK2602) was added and mixed. The sample was placed in a Thermocycler with a heated lid using the following cycling conditions: 72 °C for 5 min; 98 °C for 30 s; 14 cycles of 98 °C for 10 s, 63 °C for 30 s and 72 °C for 15s; final extension at 72 °C for 1 min and hold at 8 °C. Post-PCR clean-up was performed by adding 1.3× volume of Ampure XP beads (Beckman Counter), and libraries were incubated with beads for 5 min at RT, washed twice gently in 80% ethanol, and eluted in 30 μL 10 mM Tris pH 8.0. the library was analyzed using Agilent 2100 (2100 Bioanalyzer Instrument, G2939BA). Then the library was sequenced in MGI2000 platform with PE100+100+10 sequencing.

### BIND&MODIFY library preparation

The BIND&MODIFY library was prepared following the manufacturer’s protocol of SQK-LSK109 (Nanopore, SQK-LSK109). The library was sequenced in the ONT PromethION platform with R9.4.1 flow cell.

### Basecalling and DNA methylation calling

Reads from the ONT data were performed using megalodon (V2.2.9), which used Guppy basecaller to basecalling and Guppy model config res_dna_r941_min_modbases-all-context_v001.cfg has been released into the Rerio repository was used to identify DNA m^6^A methylation. megalodon_extras was used to get per read modified_bases from the megalodon basecalls and mappings results. In order to futher explore the accurate threshold of methylation probability, a control sample with almost no m^6^A methylation was used as background noise, and Gaussian mixture model was used to fit the methylation probability distribution generated by megalodon.

### Accessibility score

Hg19 genome and the gene elements were processed into 50bp bin sliding 5bp by Bedtools (v2.27.1)^63^. The accessibility score over multi base-pair windows were calculated as methylation ratio = m^6^A bases in all covered reads under bin/ adenosine bases in all covered reads under bin. And the accessibility score of each single molecule in the bin was also calculated.

### ChIP-seq data processing

Demultipexed fastq files were mapped to the hg19 genome using Bowtie2(2.4.1) ^64^with the following settings: bowtie2 --end-to-end --very-sensitive --no-mixed --no-discordant --phred33 -I 10 -X 700. peaks were called using MACS2 (v.2.1.0) ^65^ with the following settings: -g 12000000-f BAMPE.

### CUT&TAG data processing

Demultipexed fastq files were mapped to the hg19 genome using Bowtie2 with the following settings: bowtie2 --end-to-end --very-sensitive --no-mixed --no-discordant -- phred33 -I 10 -X 700. Because of a constant amount of pATn5 is added to CUT&TAG reactions and brings along a fixed amount of E. coli DNA, we used bowtie2 with the parameters mentioned above to remove E. coli DNA and conduct normalization according to the CUT&TAG tutorial ^25^. CUT&TAG peaks were called using SEACR(1.3)^66^ with default parameters.

### RNA-seq analysis

RNA-seq expected counts of the MCF-7 cell lines in all replicates were corrected to be TPM, the mean TPM of all replicates was used as the expression level of each gene for subsequent analysis.

### Gene single molecular diversity

The accessibility of a single molecule in each gene was calculated, and the value in each gene was sorted from small to large by the gene unit. Then the hierarchical clustering was performed for the diversity of single molecule accessibility of each gene. KOBAS3.0 ^67^ was used for KEGG and GO analysis for each cluster.

### Assess replicate reproducibility

To study the reproducibility between replicates, the genome is split into 50 bp bins and sliding 5bp, then a Pearson correlation of the log2-transformed values of m^6^A methylation ratio in each bin is calculated between replicate datasets.

### SV calling

We used NGMLR (v0.2.7) ^68^ to compare the read of ONT to the human reference genome of hg19 to get the BAM file for comparison. Then we used samtools (v1.2) to sort the bam files.The sniffles (v1.0.12) ^68^ with the parameter --genotype -T 8 -S 8 were then used to call the structural variation on the bam file created in the previous step.

### Co-accessibility assessment

To evaluate co-accessibility patterns along the genome, we applied COA as follows. Each chromosome in the genome was split into windows of size w. For each such window (i, i + w), we identified another window (j,j+w) such that the span (i,j,w) was covered by ≥N reads. For each single spanning molecule k, accessibility scores (A) in each bin were then aggregated and binarized as described above. The local co-accessibility matrix between two windows was calculated:

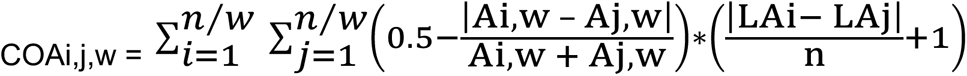

where n is the length of selected region, L is the location of region.

### Data availability

CUT&TAG for H3K27me3 and CTCF as well as Nanopore raw data are available at China National GeneBank (CNGB) with project number of CNP0001299.

### External sequencing datasets

A number of previously published MCF-7 breast cancer datasets were used in this study. ChIP-seq data for CTCF was downloaded from ENCODE with accession ENCSR000DMR, ChIP-seq data for H3K27me3 was downloaded from ENCODE with accession ENCSR761DLU. The RNA-seq data of MCF-7 was download from the Gene Expression Omnibus (GEO) repository database with the accession number GSE71862.

### Other external datasets

Hg19 genome, short interspersed nuclear elements (SINE) and long interspersed nuclear elements (LINE) region were downloaded from NCBI. TES, TTS and other gene elements were downloaded from the UCSC Table Browser. MCF-7 CTCF binding site was downloaded from the CTCFBSDB_v2.0 ^69^.

## Acknowledgment

This research was supported by the Science, Technology, and Innovation Commission of Shenzhen Municipality (grant number JSGG20170824152728492). The supporter had no role in designing the study, data collection, analysis, and interpretation, or in writing the manuscript.

## Author contributions

CT designed and supervised the experiments. ZW and FR perform the lab experiments; WTC and CT perform the bioinformatics data analysis. All others joined the data analysis.

