## Supplementary figures for "Long-range single-molecule mapping of chromatin modification in eukaryotes"

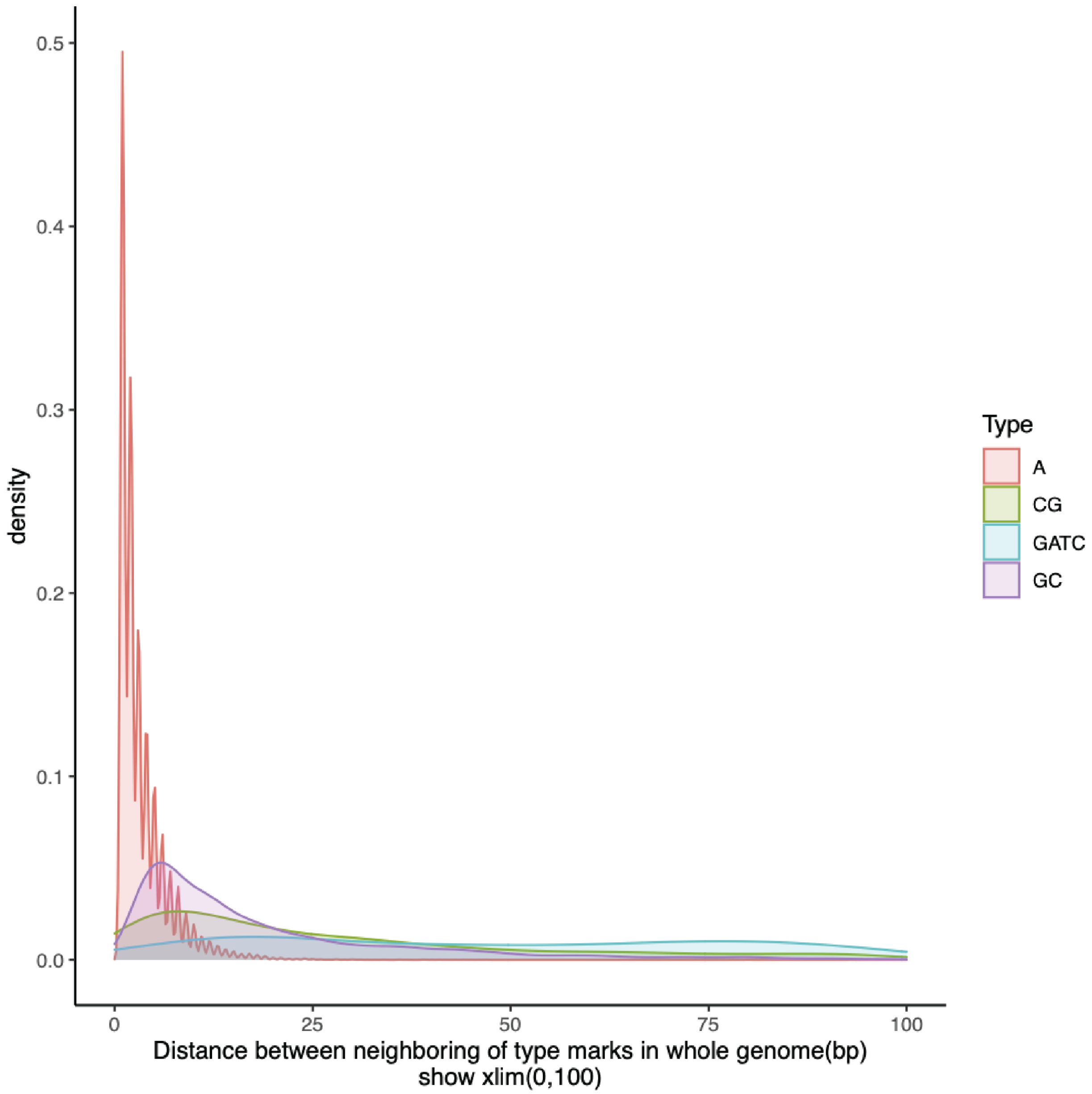


**Supplemental Figure 1. The frequency of the motif in genome.**

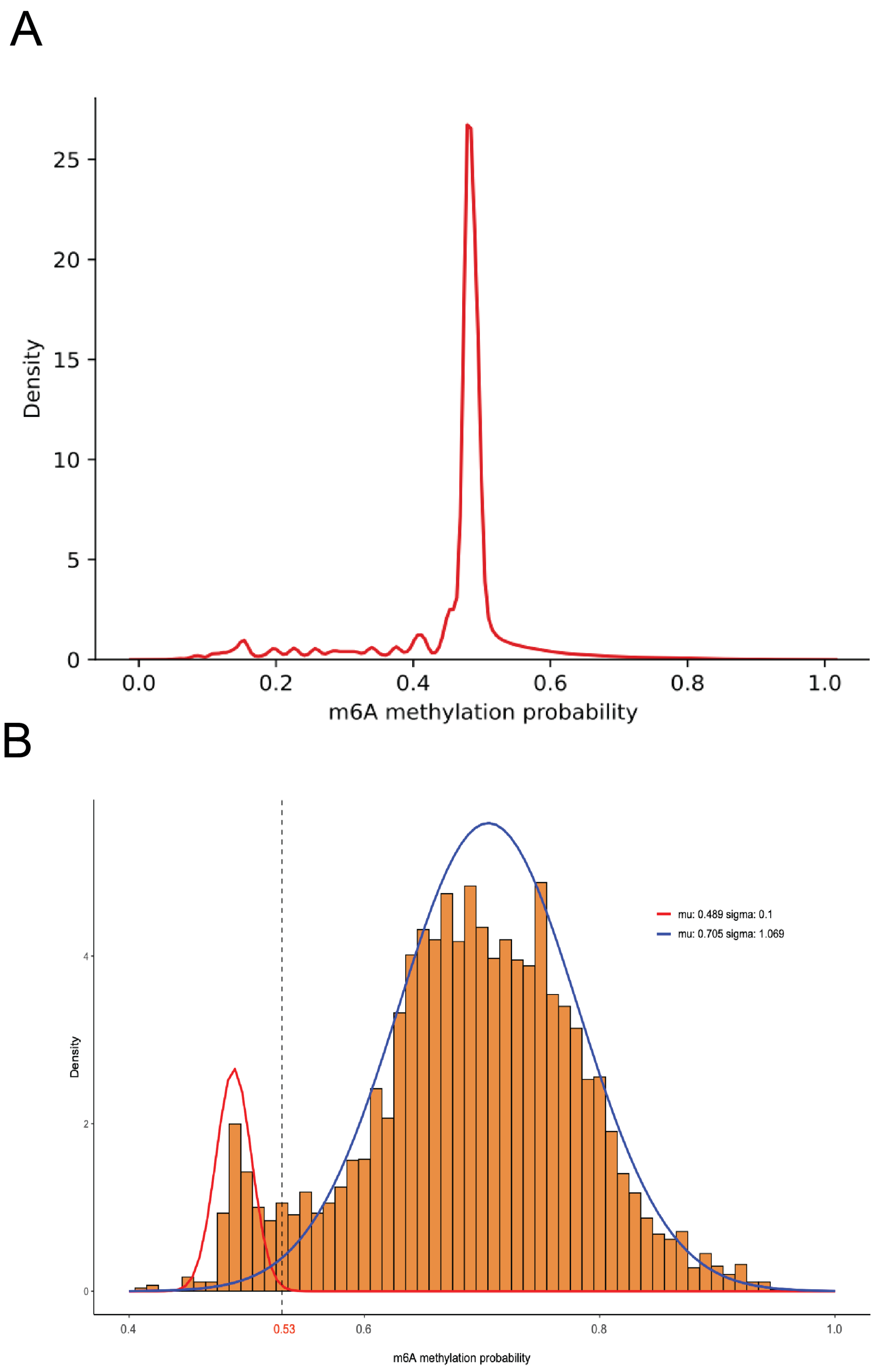
A
Density
25
20
15
10
5
0
0.0
0.2
0.4
0.6
0.8
1.0
m6A methylation probability
B
Density
4
2
0
mu: 0.489 sigma: 0.1
mu: 0.705 sigma: 1.069
0.4
0.53
0.6
0.8
1.0
m6A methylation probability

**Supplemental Figure 2. The data training of the m6A calling in the BIND&MODIFY.** (A) The density map of the m6A calling possibility in negative control. The MCF-7 cells tethered with IgG antibody and subsequently treated with pA-M.EcoGII was set as negative control and used as the background m6A noise. The x-axis was the m6A calling possibility in algorithm. (B) The density map of the m6A calling possibility in pA-M.EcoGII treated genomic DNA. By the mathematics modeling, we could find two peaks. The sharp peak represented the non-modified sites, and the wide peak represented the truly modified sites. The m6A possibility 0.53 was chosen as the cut-off value.

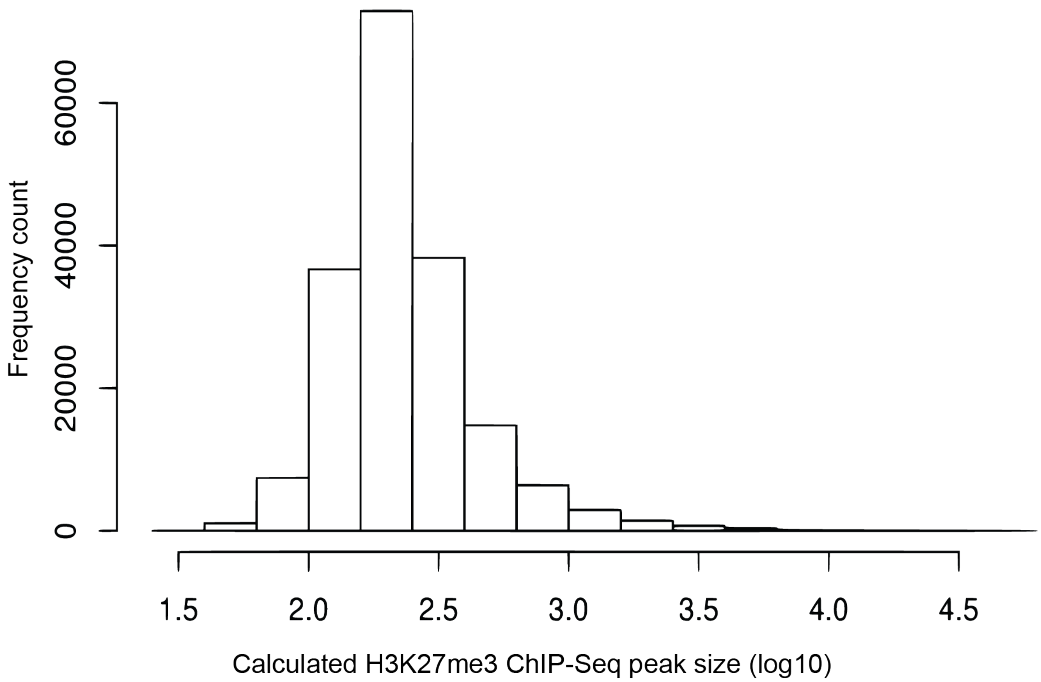


**Supplemental Figure 3. The normalized peak size distribution of the ChIP-seq.**

Most ChIP peak size distributed between 100-300bp.

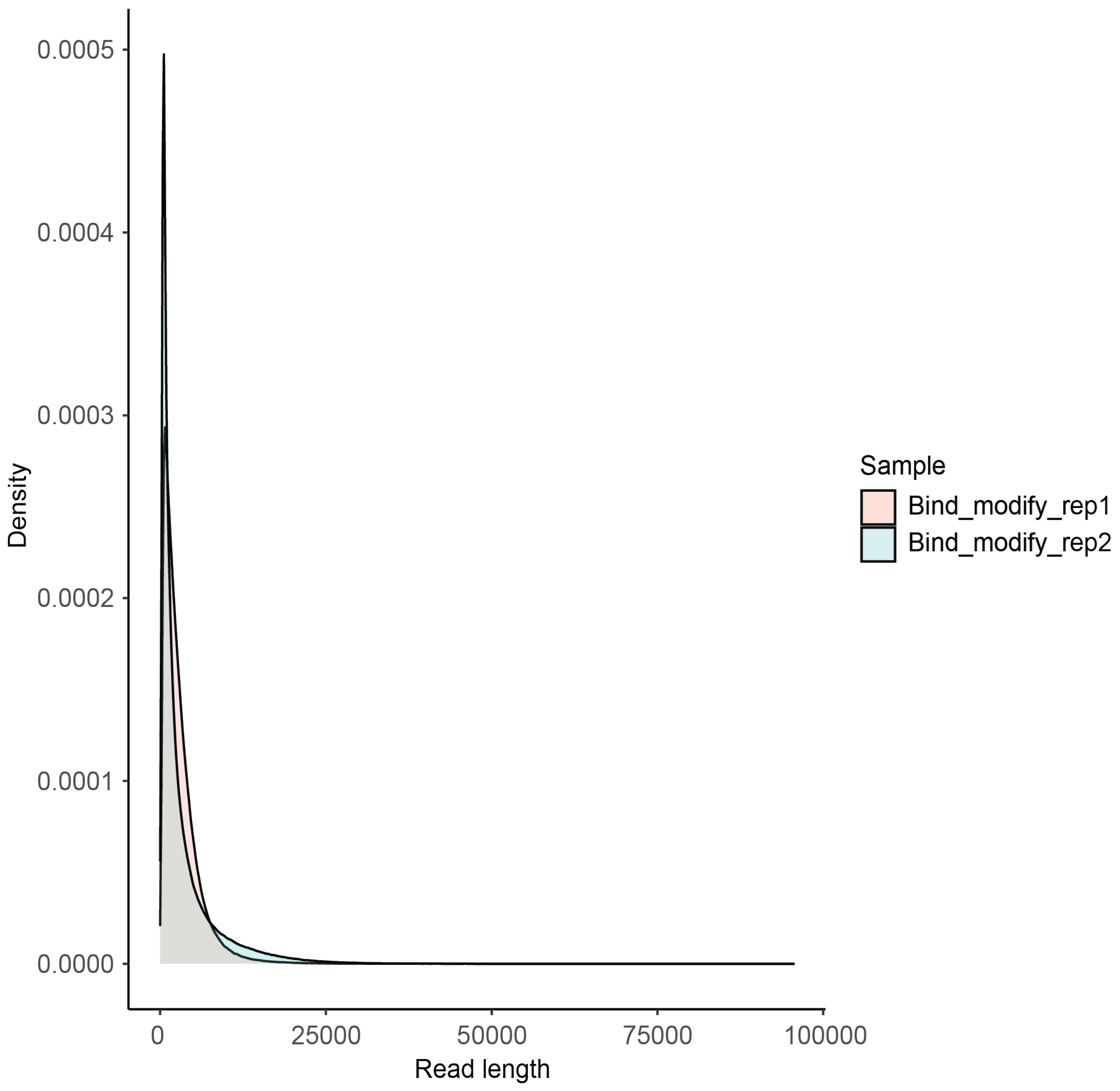


**Supplemental Figure 4. The read length distribution of the nanopore sequencing in BIND&MODIFY experiments.**

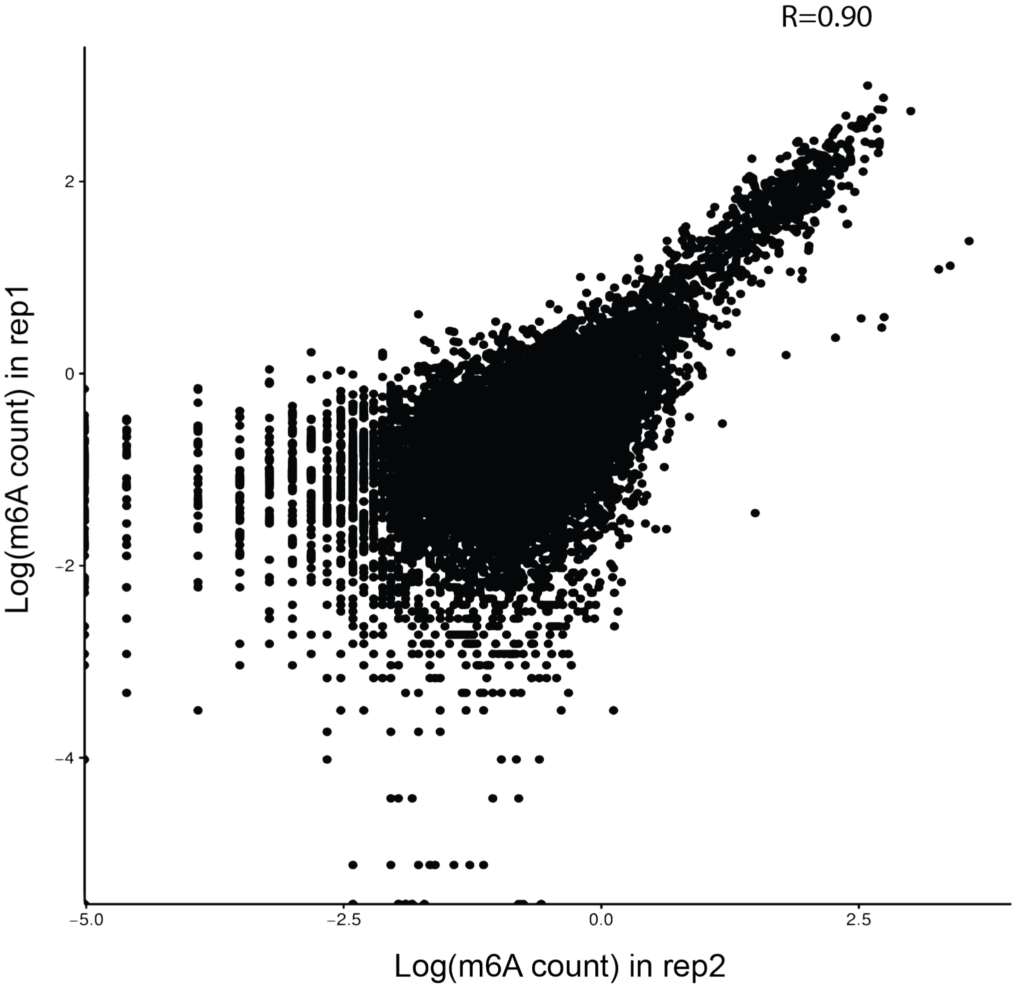


**Supplemental Figure 5. The reproducibility of the BIND&MODIFY method**. The x-axis and y-axis represented the log m6A counts of genome 50bp bins in two replicates. The two replicates (rep1 and rep2) were two technological replicates, which was done in different experimental trials (including cell culture, antibody binding, pA-M.EcoGII binding, sequencing).

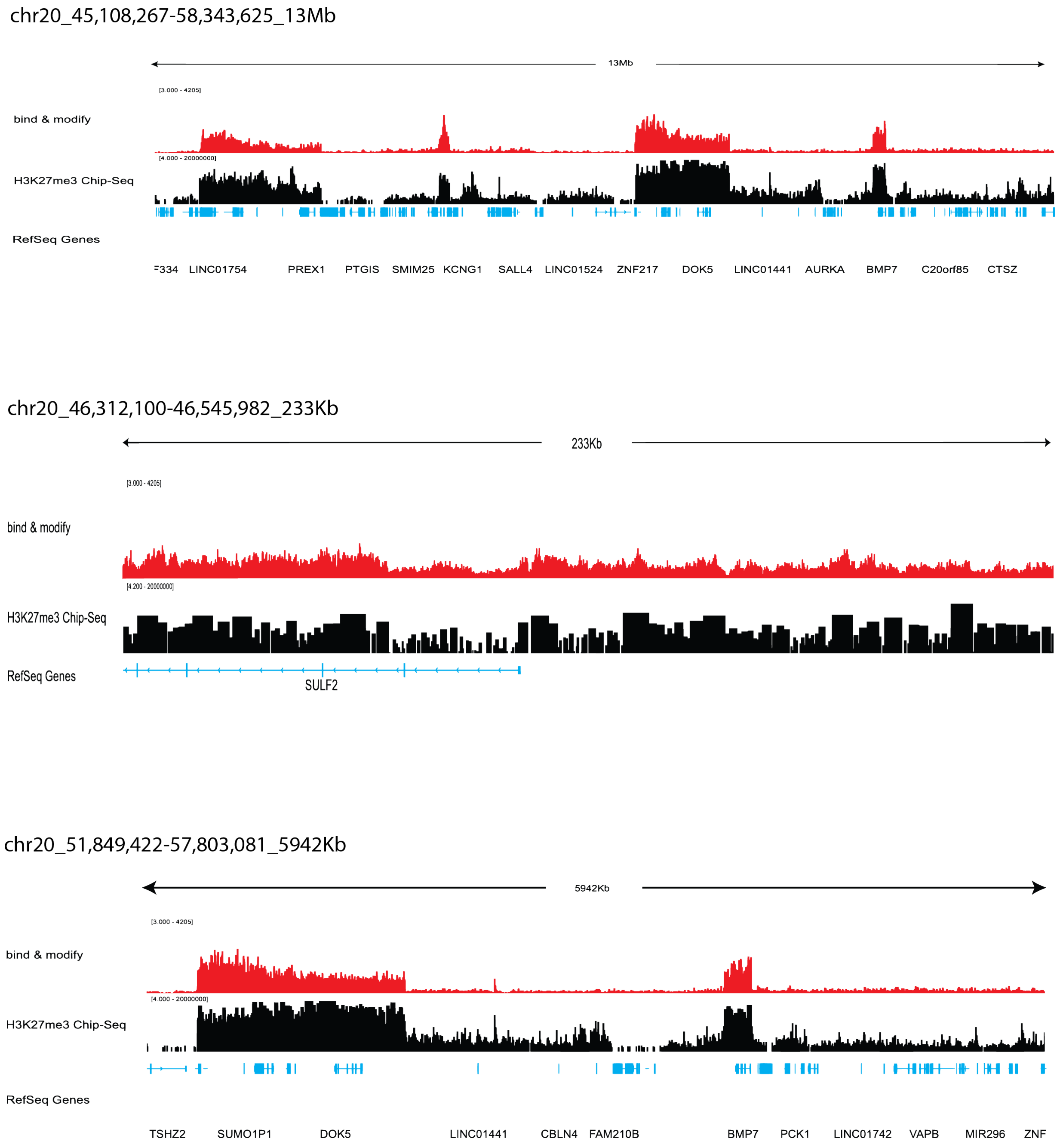


**Supplemental Figure 6. The H3K27me3 signal, by BIND&MODIFY and ChIP-seq, in genome scale view.**

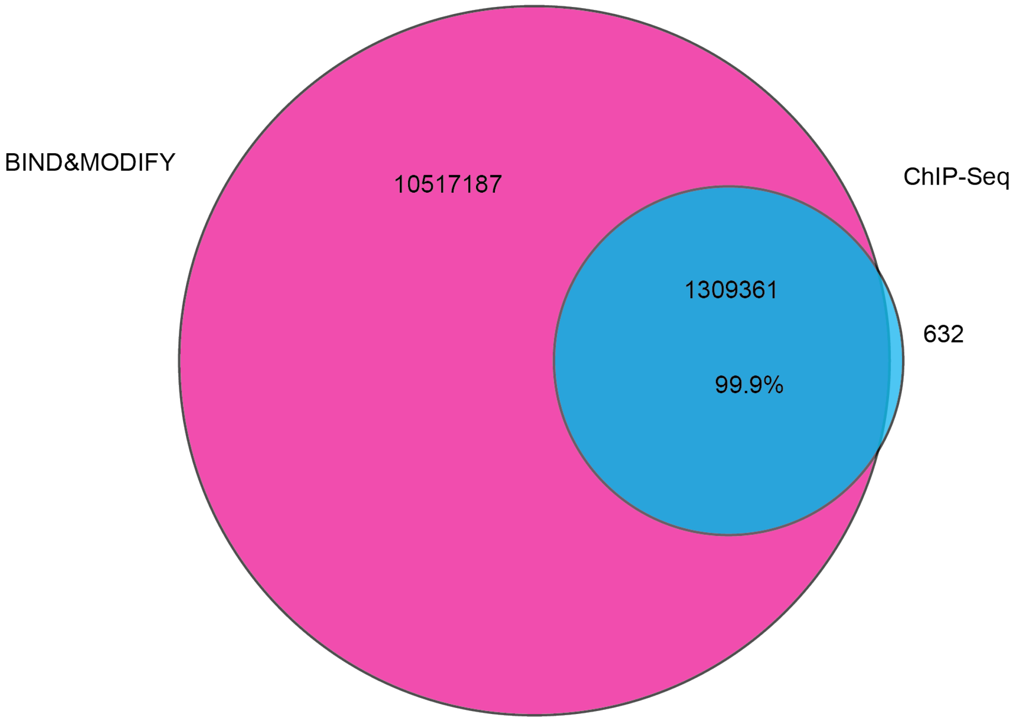


**Supplemental Figure 7. Venn diagram of high confidence position signals overlap in ChIP-seq and BIND&MODIFY.** The genome was segmentate to 50bp bins. The signal region in BIND&MODIFY was identified by any m6A counts. The signal region in ChIP-seq was identified by common peak region in two replicates.

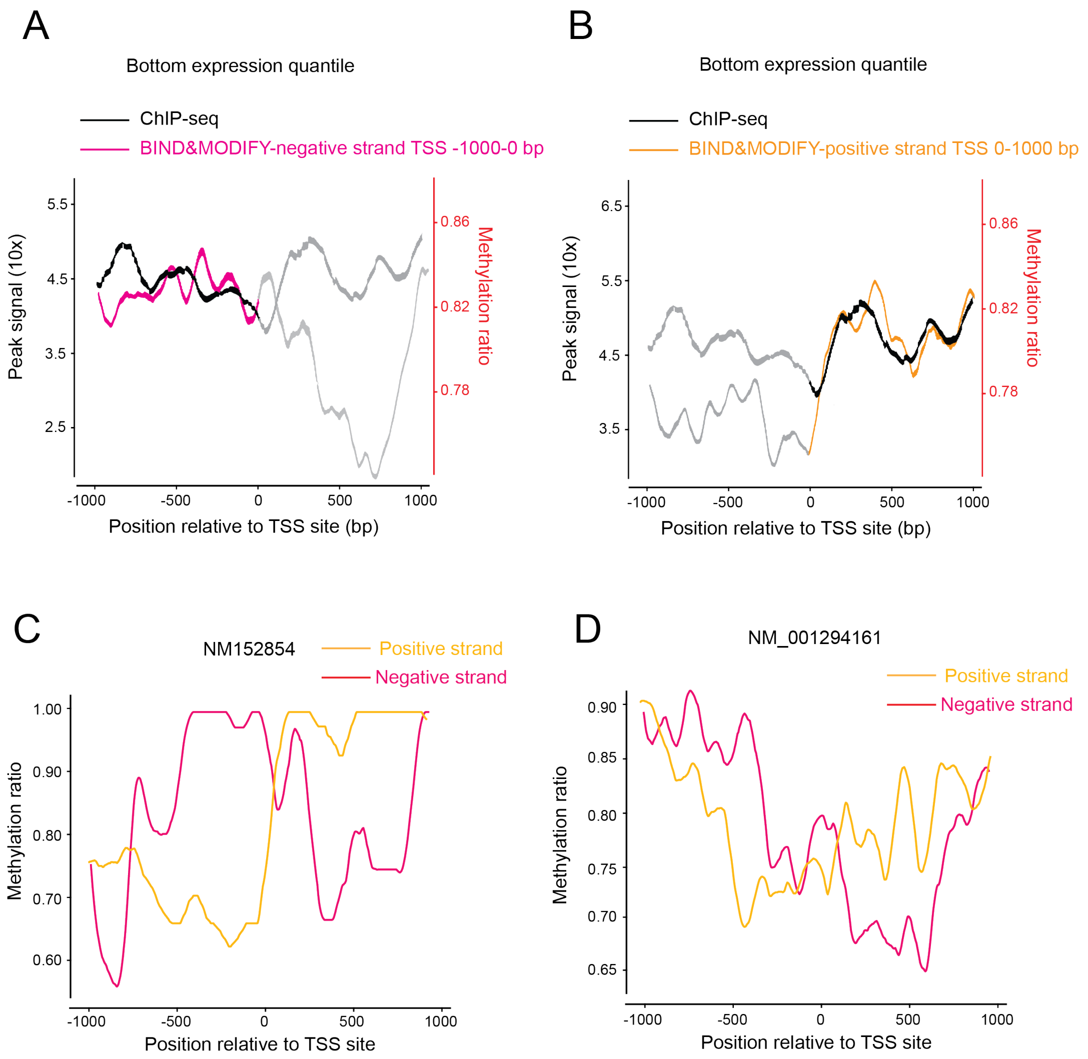


**Supplemental Figure 8. The H3K27me3 pattern in BIND&MODIFY and ChIP-seq.** (A-B) The H3K27me3 strand-specific view by the TSS centered plot for the low expression genes. The low expression genes are the genes in the bottom expression quantile. The plots covered the upstream/downstream 1000bp from TSSs. (A) ChIP-seq v.s. BIND&MODIFY negative strand-specific view. (B) ChIP-seq v.s. BIND&MODIFY positive strand-specific view. (C-D) The H3K27me3 strand-specific view by the TSS centered plot for NM_152854 and NM_001294161.

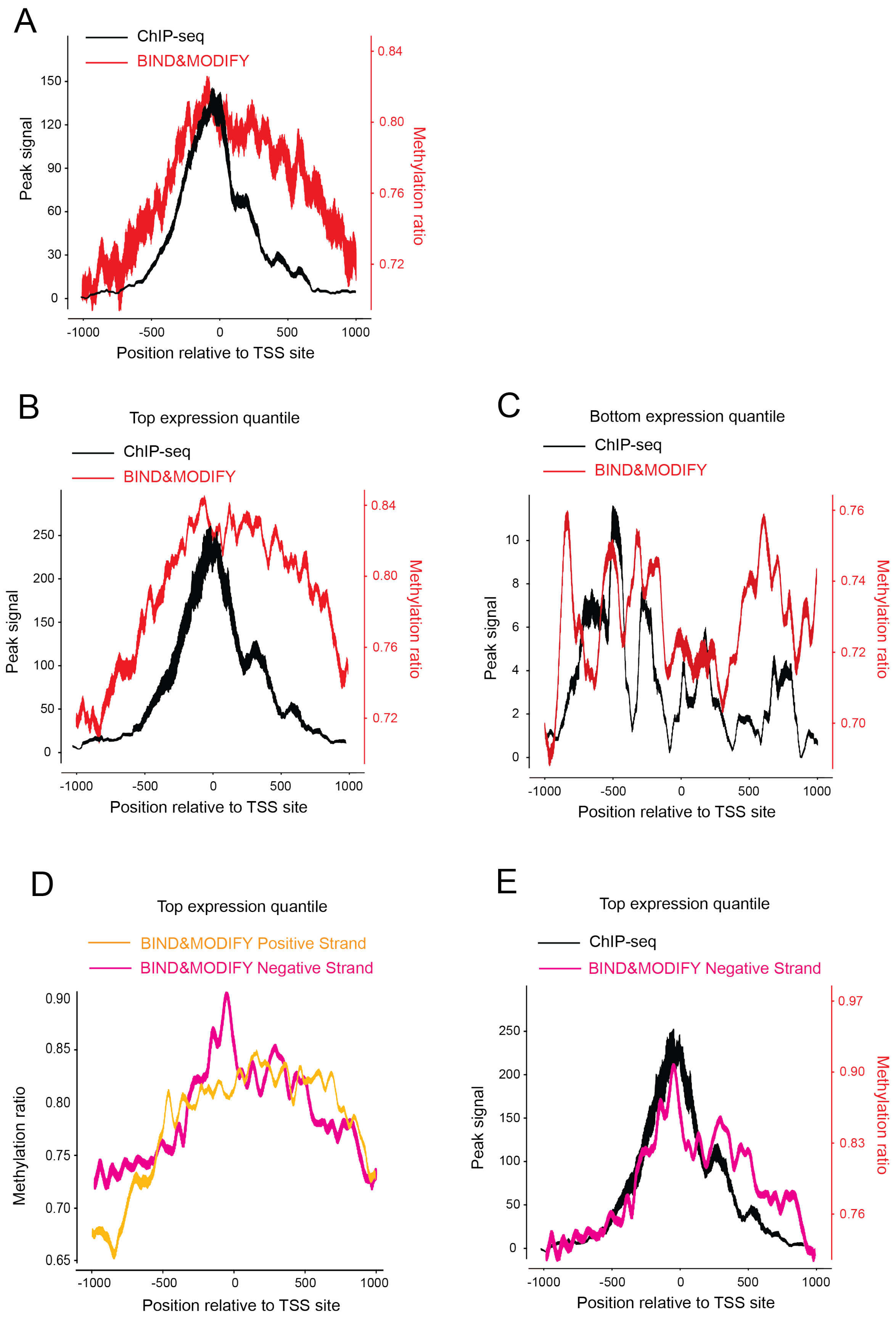
A
ChIP-seq
BIND&MODIFY
Peak signal
0
30
60
90
120
150
0.72
0.76
0.80
0.84
Methylation ratio
-1000
-500
0
500
1000
Position relative to TSS site
B
Top expression quantile
ChIP-seq
BIND&MODIFY
Peak signal
0
50
100
150
200
250
0.72
0.76
0.80
0.84
Methylation ratio
-1000
-500
0
500
1000
Position relative to TSS site
C
Bottom expression quantile
ChIP-seq
BIND&MODIFY
Peak signal
0
2
4
6
8
10
0.70
0.72
0.74
0.76
Methylation ratio
-1000
-500
0
500
1000
Position relative to TSS site
D
Top expression quantile
BIND&MODIFY Positive Strand
BIND&MODIFY Negative Strand
Methylation ratio
0.65
0.70
0.75
0.80
0.85
0.90
-1000
-500
0
500
1000
Position relative to TSS site
E
Top expression quantile
ChIP-seq
BIND&MODIFY Negative Strand
Peak signal
0
50
100
150
200
250
0.76
0.83
0.90
0.97
Methylation ratio
-1000
-500
0
500
1000
Position relative to TSS site

**Supplemental Figure 9. The CTCF pattern in BIND&MODIFY and CUT&TAG.** (A) The CTCF pattern was plotted by the TSS centered plot for all the genes. The plot covered the upstream/downstream 1000bp from TSS. (B) The CTCF pattern was plotted by the TSS centered plot for the high expression genes. The high expression genes are the genes in the top expression quantile. (C) The CTCF pattern was plotted by the TSS centered plot for the low expression genes. The low expression genes are the genes in the bottom expression quantile. (D) The CTCF strand-specific view by the TSS centered plot for the high expression genes. (E) CTCF ChIP-seq pattern and BIND&MODIFY negative strand view by the TSS centered plot for the high expression genes.

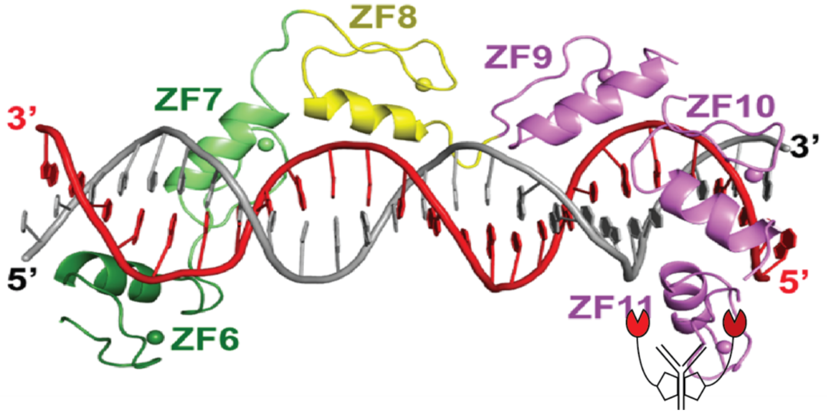


**Supplemental Figure 10. The crystal structure of CTCF binding DNA.** The antibody bound the C-terminal of CTCF, which is spatially close to the negative strand (gray).

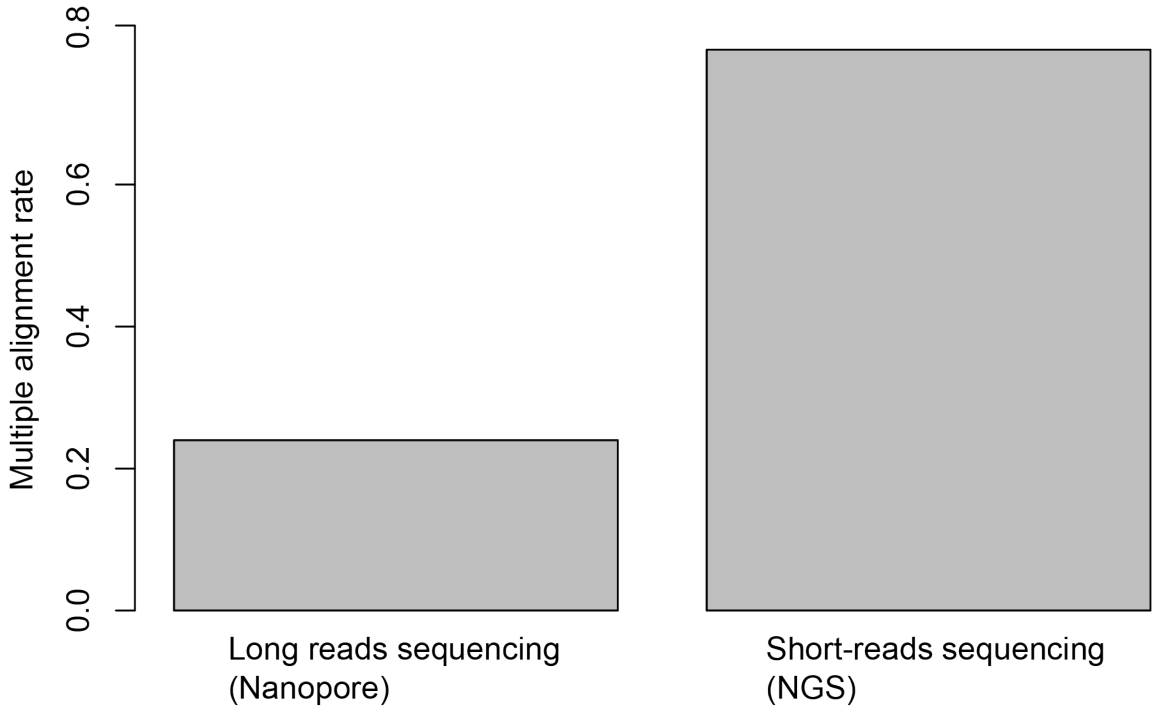


**Supplemental Figure 11. The multiple alignment rate of the ChIP-seq and BIND&MODIFY on transposon areas.**

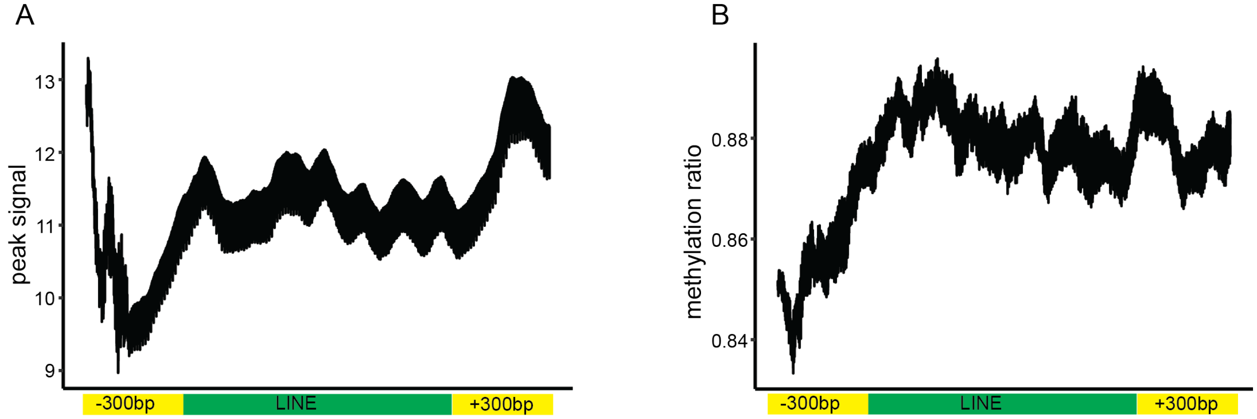


**Supplemental Figure 12. The BIND&MODIFY/ChIP-seq showed H3K27m3 pattern in LINEs.** (A-B) The LINEs with size 900-1100bp were selected and centered. The moving average H3K27me3 signals on the upstream/downstream 300bp of these LINEs, including LINEs, were plotted with corresponding genomic sites. The y-axis (A) indicated the normalized read counts of ChIP-seq. The y-axis (B) indicated the normalized methylation ratio of m6A in BIND&MODIFY with nanopore sequencing.

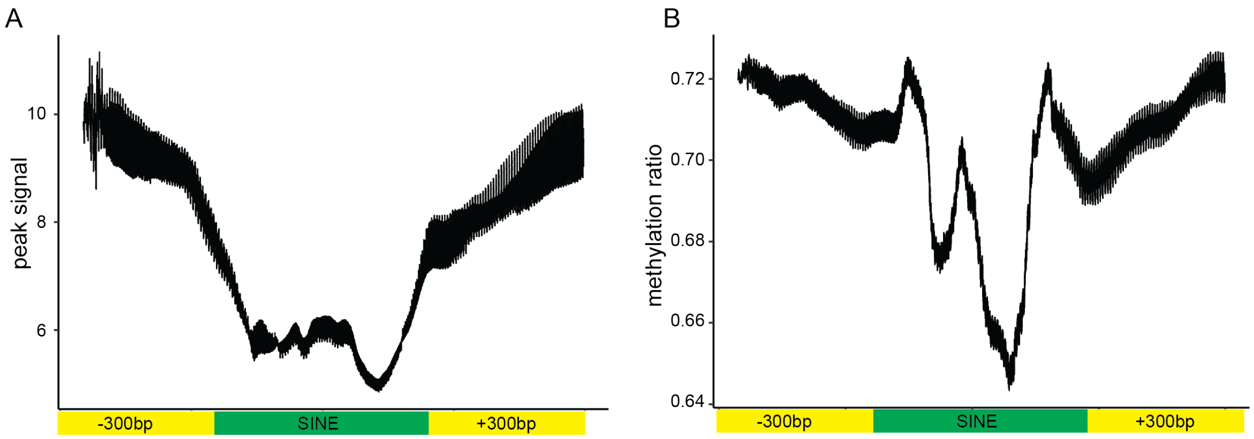


**Supplemental Figure 13. The BIND&MODIFY/ CUT&TAG showed CTCF pattern in SINEs.** (A-B) The SINEs with size 350~450bp were selected and centered. The moving average CTCF signals on the upstream/downstream 300bp of these SINEs, including SINEs, were plotted with corresponding genomic sites. They y-axis (A) indicated the normalized read counts of CUT&TAG. The y-axis (B) indicated the normalized the methylation ratio of m6A in BIND&MODIFY with nanopore sequencing.

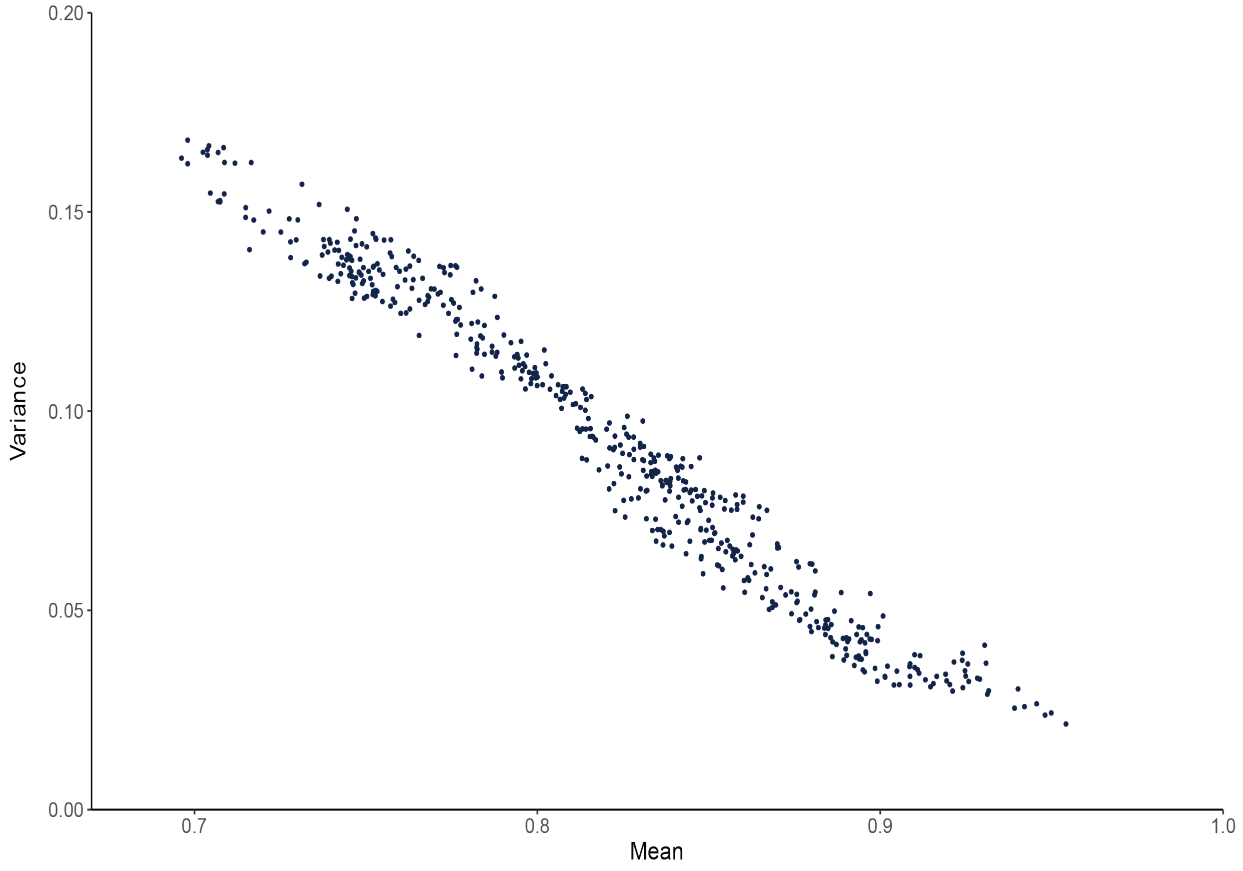


**Supplemental Figure 14. The correlation between methylation means and variance.** In the high methylation region (possibly pA-EcoGII bound region), the variance is small.

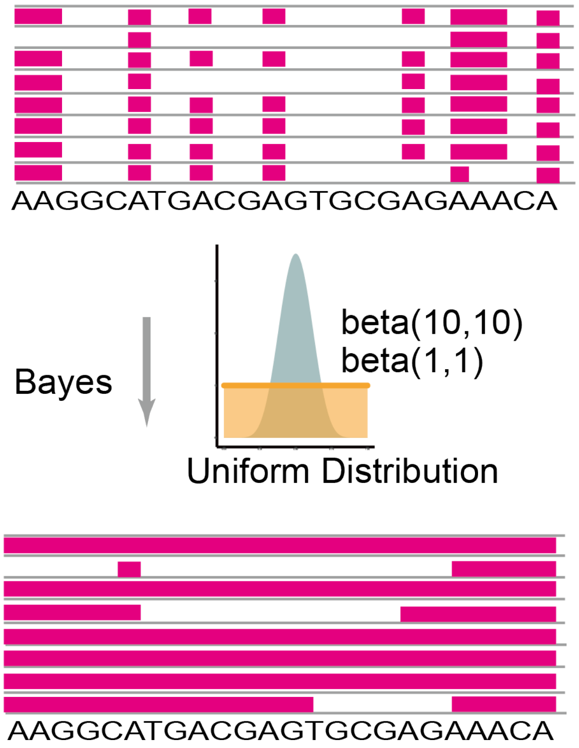


**Supplemental Figure 15. The Bayesian model calculated accumulated possibility in regions.** The methylation possibility in regions were configured with Bayesian accumulation model.

Chr20: 52,278,000~52,280,500 (GRCh37/hg19 by Entrez Gene)
Gene desert region (postive strand)

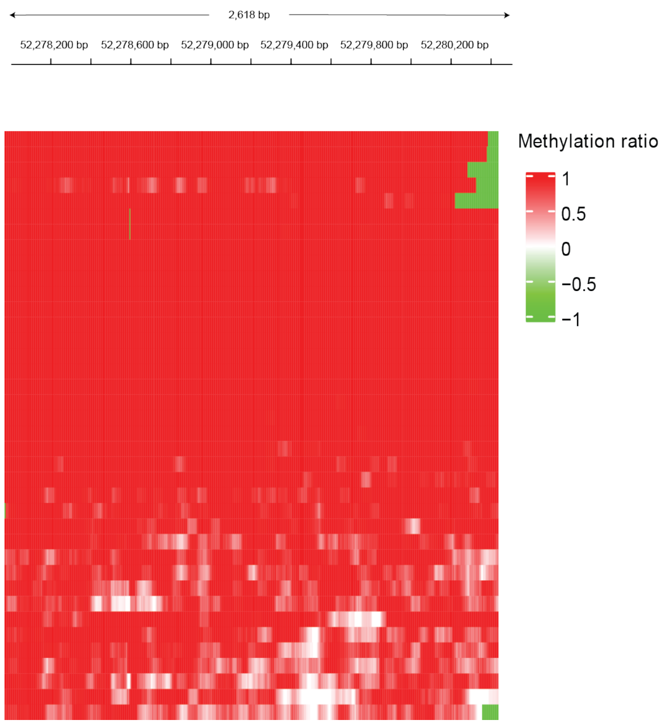


**Supplemental Figure 16. The single molecular resolution of the gene desert region Chr20: 52,278,000~52,280,500 visualize each molecular methylation statues of H3K27me3.**

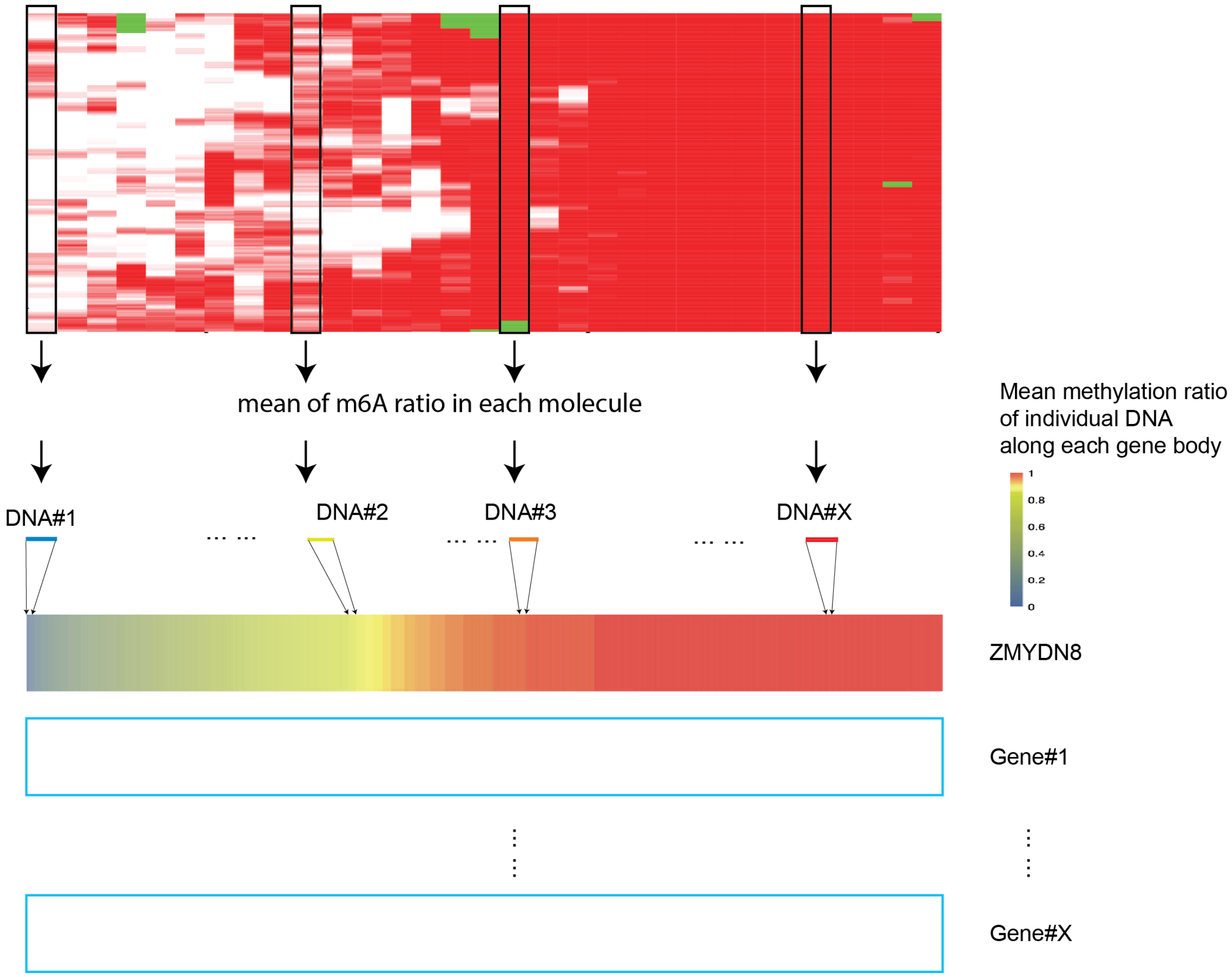


**Supplemental Figure 17. Illustration of single molecule DNA heterogeneity clustering method. (Figure 5C).**

The upper heatmap indicated the molecular heterogeneity for the genome region (90 degree of Figure 5B). Each black square box indicated one molecule. Mean methylation ratio was calculated for all the single molecule DNA that covered along each gene body, and subsequently all DNA molecules were ranked based on their mean methylation ratios for each gene. Hierarchical clustering of genes based on single molecular DNA heterogeneity was performed for the diversity of single molecule accessibility of each gene.

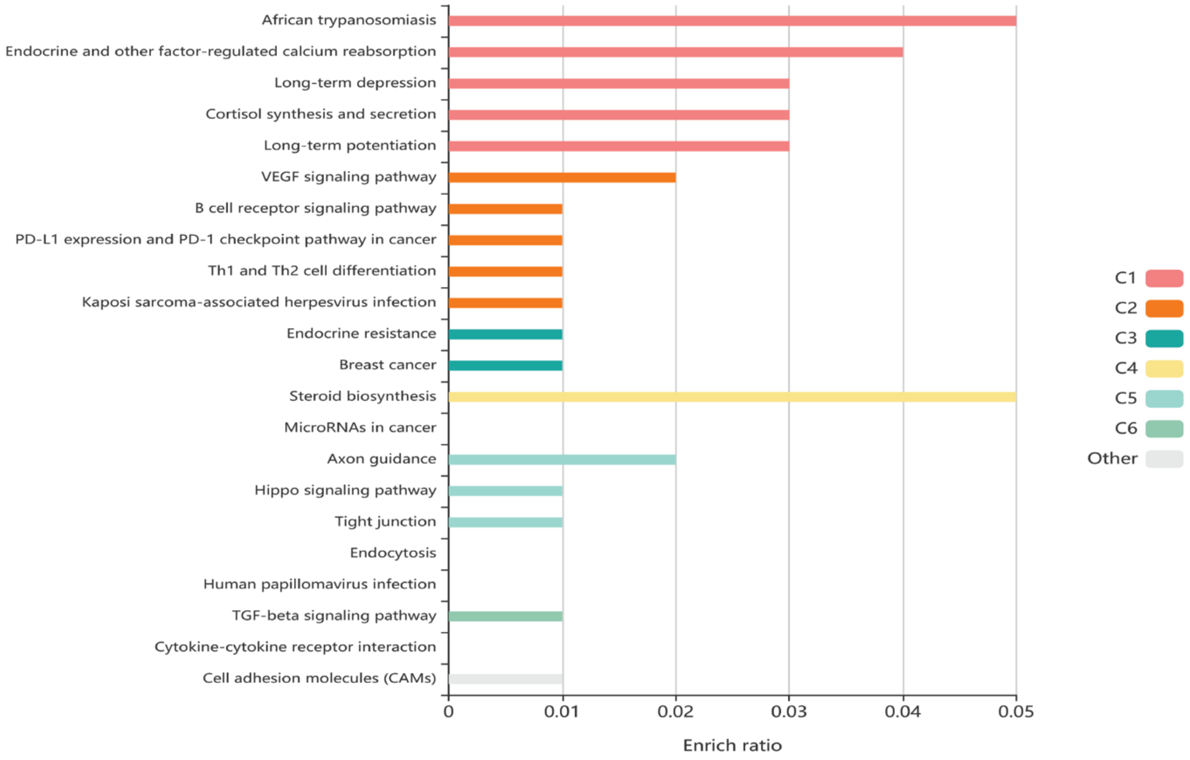


**Supplemental Figure 18. The gene ontology analysis and pathway analysis of the genes in cluster 3.** The genes in cluster 3 (Figure 5C) represent the highly heterogenous genes in chr20.

chr20_52223000-52225500.
(postive strand)

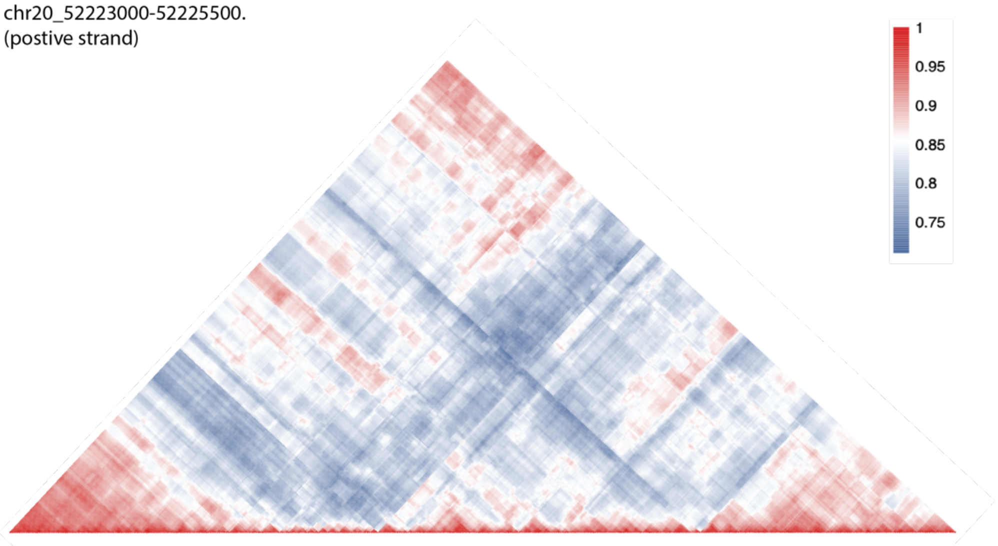


chr20_52223000-52225500.
(negative strand)

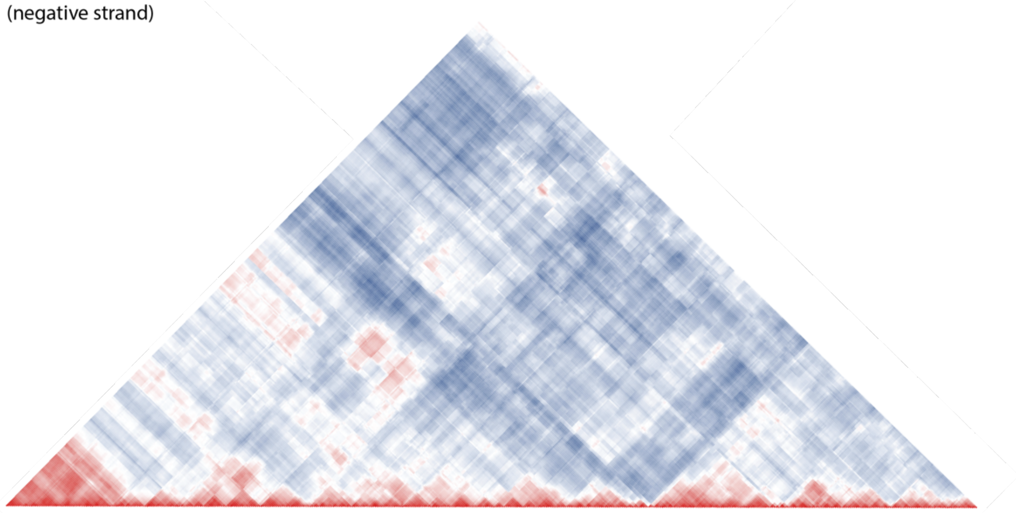

**Supplemental Figure 19**. The distance correlation of the H3K27me3 in chr20: 52223000-52225500. The upper panel is the positive strand and the lower panel is the negative strand.

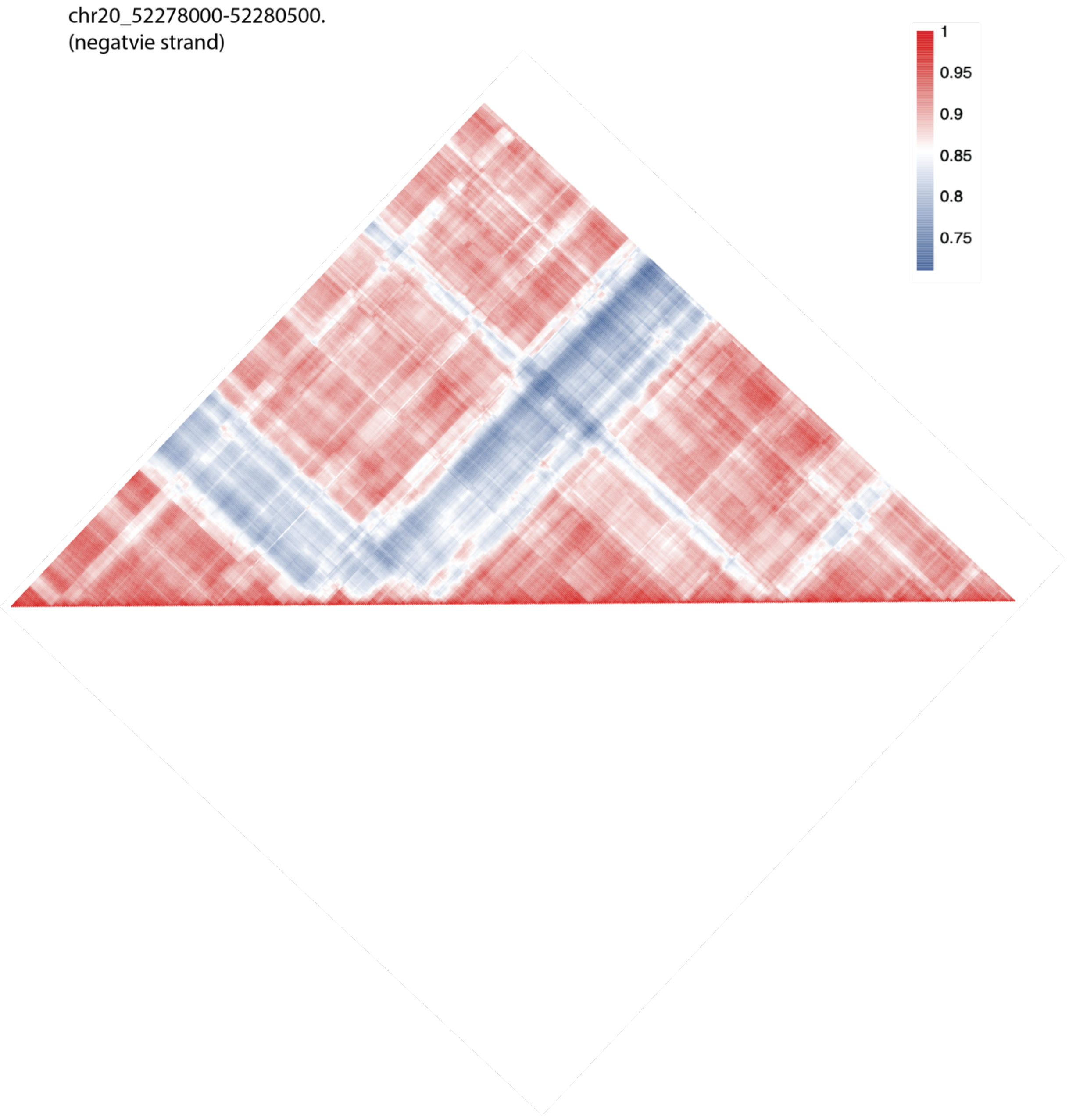


**Supplemental Figure 20**. The distance correlation of the H3K27me3 in chr20: 52278000~52280500 (negative strand).

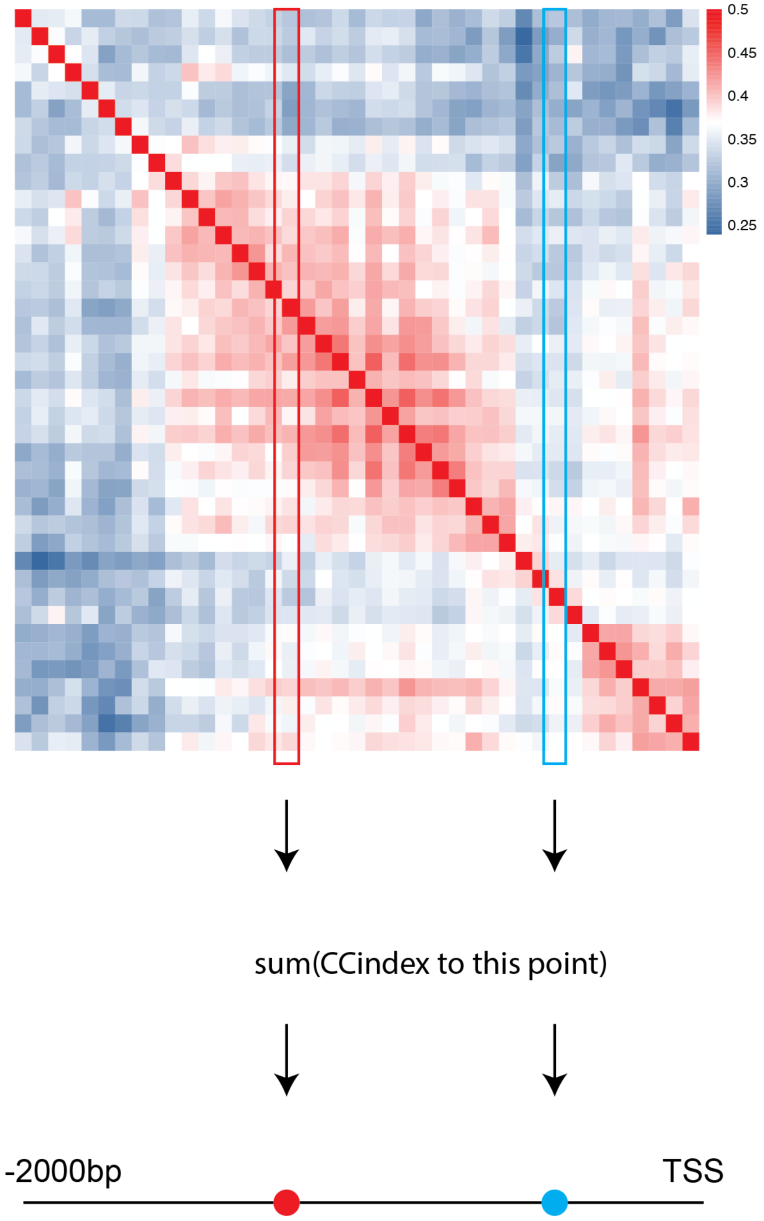


**Supplemental Figure 21. The method to calculate the Distance Effect index (DE index).** We constructed the matrix, in which each cell indicated the correlation coefficient between two corresponding locations. The upper heatmap showed the CC (correlation coefficient) indexes of the constructed matrix. The blue/red square indicate the correlation between this location and other neighboring regions. The squared cells on this location were sum up to suggested the impact of this location to other neighboring regions.

H3K27me3, Cluster 4

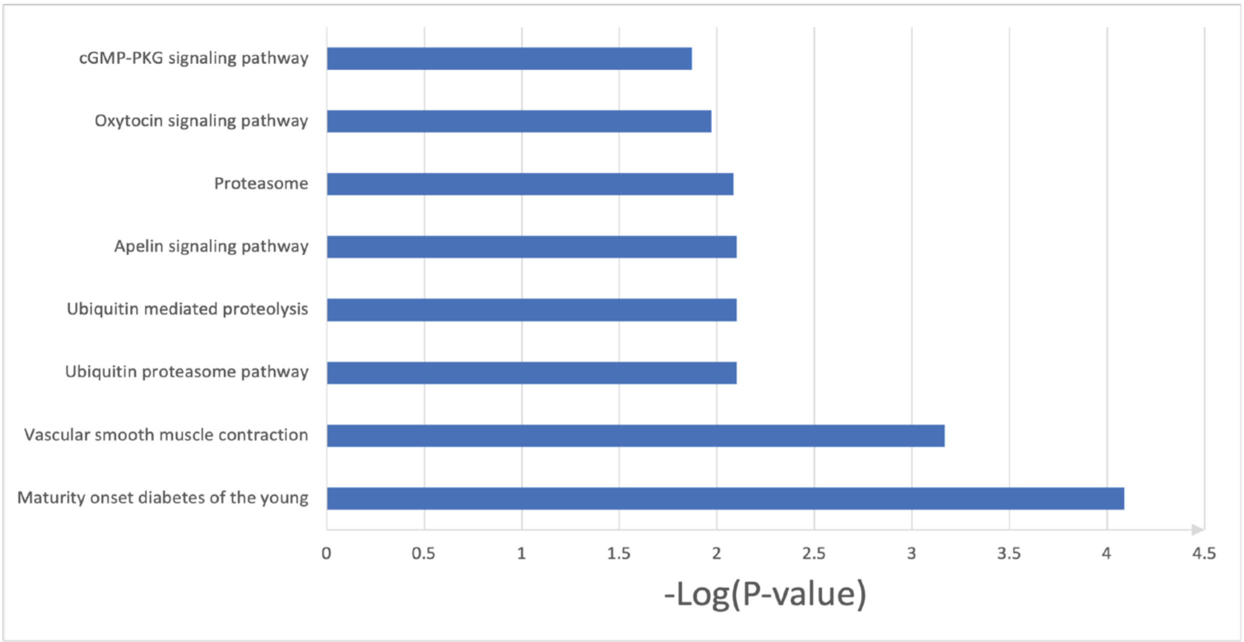


**Supplemental Figure 22**. The gene ontology analysis of the cluster 4 (Figure 5B).

CTCF, Cluster 3

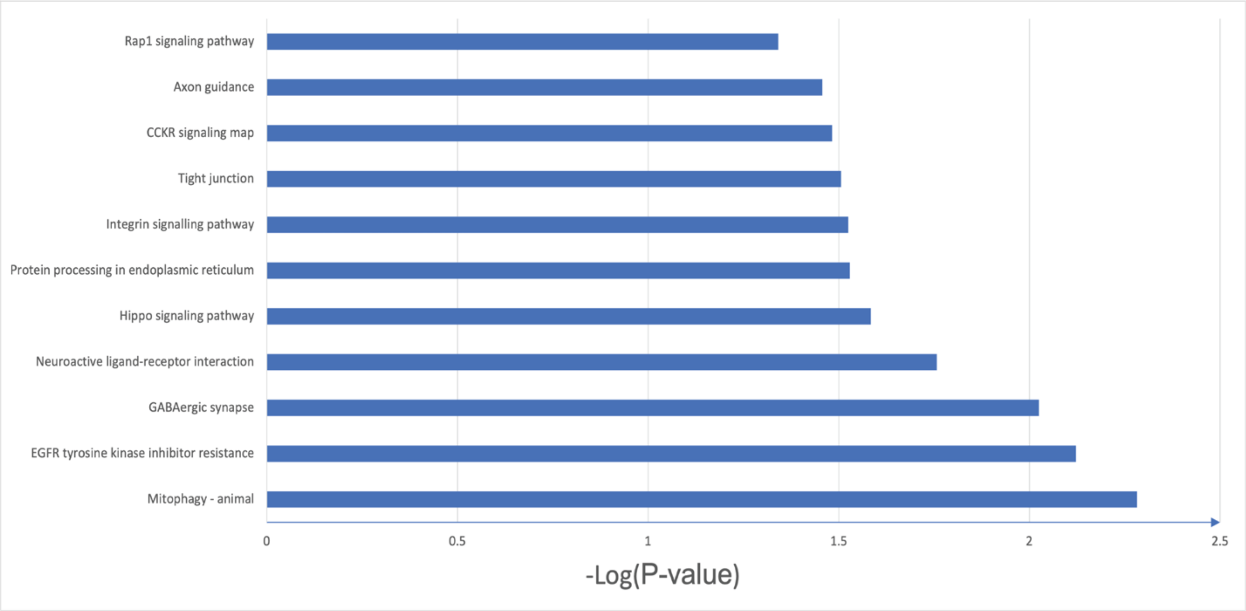


**Supplemental Figure 23.** The gene ontology analysis of the cluster 3 (Figure 5C).

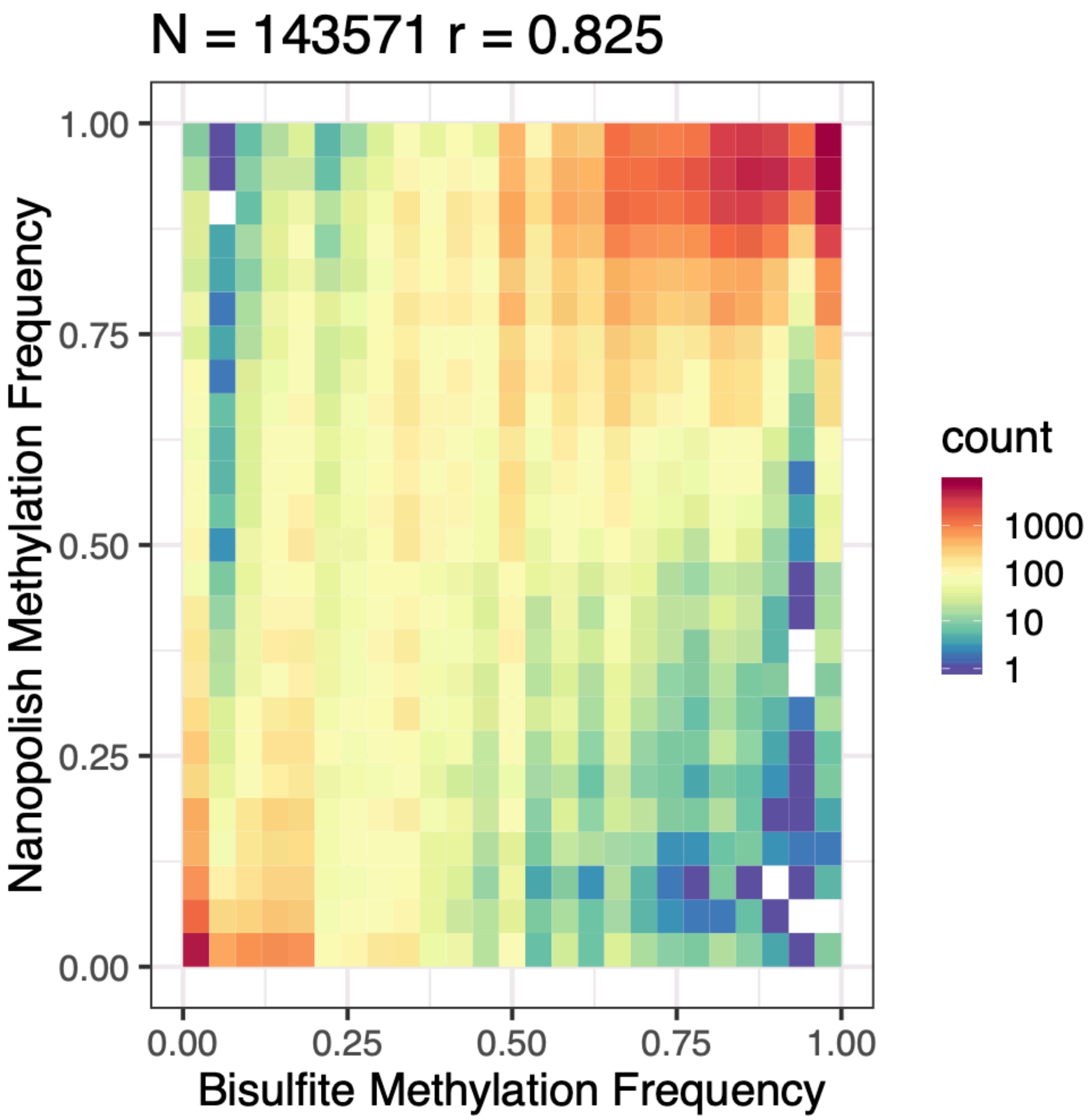


**Supplemental Figure 24**. The 5mC correlation between Nanopolish base calling and bisulfite sequencing. The correlation showed 0.83, which indicates our BIND&MODIFY m6A methylation did not influence native 5mC methylation base calling.
